# Pleistocene sea-level fluctuation shapes archipelago-wide population structure in the Endangered Lord Howe Island cockroach *Panesthia lata*

**DOI:** 10.1101/2024.11.12.623290

**Authors:** Maxim W.D. Adams, Kyle M. Ewart, Nicholas Carlile, Harley A. Rose, James A. Walker, Ian Hutton, Simon Y.W. Ho, Nathan Lo

## Abstract

Studies of biogeographic processes have often centred islands as model systems, yet questions remain about the role of Pleistocene sea-level fluctuations in shaping islands’ biodiversity. One novel, potentially informative model system is the Lord Howe Island Group of Australia. Despite the World Heritage status of this archipelago, almost nothing is known of the biogeographic origins, evolutionary distinctiveness or genetic diversity of the ecological communities across its 28 islands. In this study, we focused on the cockroach *Panesthia lata*, an ecologically specialized invertebrate with one of the broadest recorded distributions of any LHIG species. To investigate the influence of Pleistocene sea-level fluctuations on LHIG fauna, we explored the phylogeography of *P. lata* using single-nucleotide polymorphisms and complete mitochondrial genomes. Our analyses reveal that the lineage on the permanently isolated islet Ball’s Pyramid is highly divergent from the remaining populations, while those on the episodically connected Lord Howe, Roach and Blackburn Islands experienced gene flow during the last glacial period. These results offer the first evidence that Pleistocene land bridges allowed for overland migration across the archipelago. Further, although *P. lata* was believed to have been locally extirpated by rodents on Lord Howe Island, we discovered two surviving, relict populations. We also detected high levels of inbreeding in all populations, emphasizing the need for ongoing conservation management. Finally, the combination of shallow genetic structure and low diversity suggests that genetic rescue from another island may be a viable strategy to conserve the Lord Howe Island population of *P. lata*, as well as other species that have been similarly impacted by rodents.

## 1. Introduction

Islands have long been used as natural laboratories in which to study evolutionary processes, such as colonization, extinction and adaptative radiation. In recent decades, there has been increasing interest in the role of Pleistocene sea-level fluctuation in the evolution of archipelagic biotas across the globe (reviewed by Ávila et al., 2019). During ancient periods of lower sea level (“lowstands”), many presently isolated islands amalgamated into larger “palaeo-islands”, permitting overland migration and gene flow. As a result, a paradigmatic view is that contemporary populations inhabiting fragments of a single palaeo-island should be less evolutionarily distinct than those divided across permanently separated land masses, with the depth of genetic structure corresponding to the most recent lowstand (terminating *ca*. 10 ka; e.g., Barker et al., 2012; Fattorini, 2010; Fernández-Palacios et al., 2015; Papadopoulou & Knowles, 2017). However, some studies have documented archipelago-wide genetic structure that is independent of the physical connectivity of landmasses, reflecting biotic and abiotic barriers that prevented historical overland migration (e.g., Cros et al., 2020; Cros & Rheindt, 2017; Garg et al., 2018; Rijsdijk et al., 2014). Moreover, in some island systems, the episodic isolation and re-amalgamation of land masses has instead acted as a “species pump”, promoting regular bouts of speciation to produce highly divergent communities on individual fragments of palaeo-islands (Ali & Atchinson, 2014; Heaney, 1985; Li & Li, 2018; but see Papadopoulou & Knowles, 2015, 2017). The generality and relative frequency of these differing biogeographic processes remain unclear.

Most studies have focused on established model systems, such as the Malay Archipelago or Macaronesia, but further insights can be gained by examining a broader range of archipelagos (reviewed by Brown et al., 2013; Cros et al., 2020; Leonard et al., 2015; Rijsdijk et al., 2014). One such archipelago is the Lord Howe Island Group (LHIG), a UNESCO World Heritage-listed archipelago that is recognized for its exceptional natural beauty and unique biodiversity. Located *ca.* 600 km east of mainland Australia, the LHIG comprises the eroded remnants of a large shield volcano that formed *ca.* 7 million years ago (Gilmore et al., 2020). The species richness of the LHIG is unusually high for an archipelago of its size and age, with surveys documenting 241 species of vascular plant, nearly 200 birds and over 1600 terrestrial invertebrates (Cassis et al., 2003; Green, 1993; Lillemets & Wilson, 2002; McAllan et al., 2004). The LHIG is considered a major conservation prioriy by Australian state and federal authorities (DECC, 2007), and is presently managed under a Permanent Park Preserve.

Over 95% of the archipelago’s land area is represented by Lord Howe Island proper (hereafter LHI), a mountainous island covered predominantly by rainforest (Sheringham et al., 2016). The remaining terrestrial habitat is divided between 27 smaller islets, which sustain comparatively xeric ecosystems dominated by grasses and sedges (Carlile & Priddel, 2013a, 2013b, 2013c, 2013d, 2013e; Carlile et al., 2013; Figure 1). The LHIG is a potentially informative system for biogeographic study, given that almost all of its islands have been episodically connected by Pleistocene land bridges, with only Ball’s Pyramid remaining isolated over the long term (constituting two palaeo-islands; Rogers et al., 2023). However, compared with LHI, the islets have received little scientific attention; in fact, most have never been formally surveyed. In addition, there have been no studies of genetic structure across multiple islands in any species from the archipelago. Consequently, it is completely unknown whether sea-level fluctuation has resulted in episodic gene flow across the LHI palaeo-island, acted as a species pump to increase genetic differentiation across the LHI palaeo-island, or even negligibly affected patterns of genetic variation across the LHIG.

**Figure 1.**
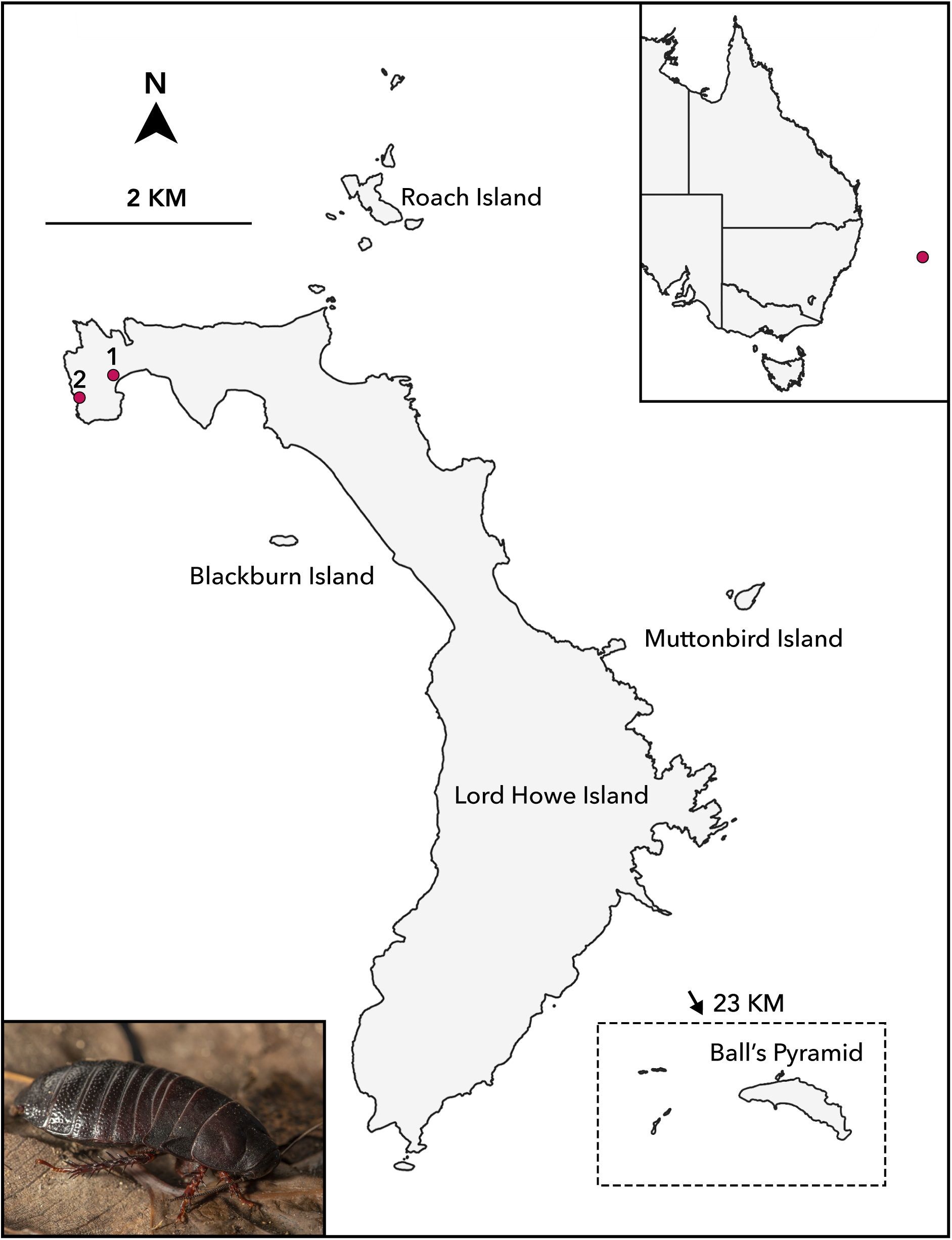
Map of the Lord Howe Island Group, showing contemporary arrangement of islands. Ball’s Pyramid is the only land mass to have remained permanently isolated throughout Pleistocene sea-level fluctuations. Markers denote two putative relict populations of *Panesthia lata* on Lord Howe Island, in North Bay (1) and North Head (2), respectively. **Top right:** Position of the islands relative to eastern Australia. **Bottom left:** *P. lata* in lateral view. Photograph: J. Gilligan.

In recent years, the question of archipelago-wide genetic structure has gained relevance for conservation of the LHIG’s species. Following human settlement in 1834, 20 exotic predators have established feral populations on LHI (Anderson, 2003). The introductions of mice (*Mus musculus*) in the 1860s and ship rats (*Rattus rattus*) in 1918 had especially large impacts, precipitating the extinctions of at least 20 endemic species, and declines of potentially hundreds more (DECC, 2007; Hutton et al., 2007; Wilkinson & Priddel, 2011). Despite almost a century of targeted management via hunting, biocontrol and poison baiting, a 2016 survey estimated that populations of rats and mice numbered up to 150,000 and 210,000, respectively (Lord Howe Island Board, 2016). In 2019, after more than a decade of planning, both species were successfully extirpated from LHI, representing the largest rodent eradication ever undertaken on an inhabited island. While many prey populations have begun to rebound, a large number remain substantially reduced and fragmented (N. Carlile, unpub. data). Importantly, all the smaller islets remained completely rodent-free throughout this period, serving as refugia for numerous birds, reptiles and invertebrates (Cassis et al., 2003; Reid et al., 2020). These now represent potentially significant insurance populations and sources of propagules for conservation translocations (e.g., DECC, 2007; Hutton et al., 2007). However, the conservation strategies for the islets will depend on their communities’ genetic distinctiveness and diversity. Therefore, a focused phylogeographic study that includes an estimate of population divergence times has the potential to illuminate the LHIG’s biogeographic history and also inform its future management.

One promising candidate for such a study is the Endangered Lord Howe Island cockroach *Panesthia lata* (Walker, 1868). A large (∼40 mm), flightless insect, *P. lata* is an ecologically important decomposer, feeding on decaying plant material and burrowing into shallow topsoil (Carlile et al., 2018). The species has one of the broadest recorded distributions of any LHIG species, spanning LHI, Ball’s Pyramid, Blackburn Island and Roach Island, and potentially other unsurveyed islets. Previous phylogenetic studies have estimated that the islet populations diverged either 2–5 Ma (Lo et al., 2016) or *ca.* 330 ka (Adams et al., 2024), potentially evidencing long-term isolation over multiple sea-level cycles. However, these studies lacked the geographic and molecular resolution to fully characterize genetic structure across the species’ range over both LHIG palaeo-islands.

Following the arrival of ship rats, observations of *P. lata* declined precipitously on LHI (reviewed by Carlile et al., 2018). In 2004, the New South Wales Scientific Committee evaluated that the cockroach had likely become locally extinct on LHI (NSW Scientific Committee, 2004), although no formal surveys have subsequently been undertaken to confirm the species’ absence. The remaining populations on Roach and Blackburn islands both appear small and spatially restricted (Carlile et al., 2018; Rose, 2003), and little is known about the species’ ecology on Ball’s Pyramid.

In this study, we undertake the first population genetic analysis across multiple islands of the LHIG, utilizing both contemporary and museum samples of *P. lata* from across the species’ known range. Our analyses allow us to address two key aims: 1) to test whether genetic structure in *P. lata* reflects the historical connectivity of palaeo-islands, or whether genetic structure is independent of ancient land bridges; and 2) to characterize the genetic health of each population and inform conservation priorities. In view of the low vagility and wide distribution of *P. lata*, we consider the species to be an ideal model for investigating the biogeographic history of the LHIG, and hope that our results lay the groundwork for the management of other organisms across the archipelago.

## 2. Methods

### 2.1. Taxon sampling and DNA extraction

Samples of *P. lata* were collected from throughout the species’ known habitat range. In August and July 2022, we retrieved specimens from Blackburn Island (*n* = 15) and Roach Island (*n* = 5). Because only a small number of samples were found on Roach Island in 2023, we also included specimens collected from this location in December 2000 (*n* = 6) and December 2003 (*n* = 11), held in the Australian Museum, Sydney, and the private collection of H.A. Rose, respectively. All samples were stored in 70–100% ethanol prior to DNA extraction.

Due to the inaccessibility of Ball’s Pyramid, we subsampled tissue from specimens collected in 1969 (*n* = 2), which represent the only known material from this population. We also subsampled specimens that had been collected on LHI between 1869 and 1950 to allow us to characterize genetic diversity prior to rodent incursion (*n* = 20). These historical samples were sourced from the pinned collections of the Macleay Museum, Sydney and the Australian Museum, Sydney (Supplementary Table S1).

Further, in July 2022, we opportunistically examined sites across LHI for the presence of *P. lata*, resulting in the discovery of a potential relict population in the island’s North Bay (Figure 1). Based on coarse estimates of the population’s density, we retrieved 10 specimens from the population, which represented considerably less than 10% of the total abundance and thereby avoided unnecessary risk to the population. In May 2023, an additional putative LHI population was discovered on the island’s North Head (Figure 1). However, the discovery occurred after the completion of DNA sequencing and this population was omitted from the present study. The discovery and biology of the putative relict populations will be discussed in a focused follow-up study (Coady et al., in prep.).

### 2.2. DNA extraction and SNP library preparation

DNA was extracted from leg muscle tissue. For contemporary specimens (< 25 years old), we extracted DNA using a DNeasy Blood and Tissue Kit (Qiagen, Germany) per the manufacturer’s instructions. The purity and concentration of each DNA sample was measured using a NanoDrop 2000 spectrophotometer (Thermo Fisher Scientific, USA) and a Qubit 2.0 fluorometer (Invitrogen, USA), respectively. For historical samples (> 25 years old), DNA extraction was undertaken at the Australian National Insect Collection, Canberra, utilizing a proteinase K digestion and silica filter-based approach that is suitable for fragmented DNA (see Jin et al., 2020; Zwick & Zwick, 2023 for methods). In total, 68 taxa were sampled for SNP generation (Blackburn Island, *n =* 15; Roach Island, *n* = 21; North Bay, *n* = 10; historical LHI, *n* = 20; Ball’s Pyramid, *n* = 2; Supplementary Table S1).

SNP data were generated by Diversity Arrays Technology (DArT), Canberra, using the DArTseq^TM^ reduced-representation sequencing platform (following methods in Cruz et al., 2013; Kilian et al., 2012). Briefly, genome complexity was reduced by digestion with the *PstI* and *HpalI* restriction enzymes, then sequenced on a NovaSeq 6000 S1 with 138-bp single-end reads. The resultant short-read sequences were processed through the DArT analytical pipeline, which removes poor-quality sequences, demultiplexes reads, and subsequently calls SNPs using the proprietary DArTsoft14 algorithms. SNPs were called against a reference genome of the closely related species *Panesthia cribrata* (Ewart et al., 2024). Due to DNA degradation, no historical LHI samples were successfully genotyped.

SNP data were filtered primarily using the R package *dartR* v.2.7.2 (Gruber et al., 2018) in RStudio (RStudio Team, 2020), under varying criteria to suit different analyses. First, we removed SNPs that were potentially erroneous or that had high levels of missing data, based on reproducibility (100%), minor allele frequency (MAF > 0.03; based on 3/2*N*) and call rate (> 0.8). Individuals with high levels of missing data (call rate < 0.1) were also removed. This initial (“quality-controlled”) data set comprised 10,370 SNPs from 48 individuals.

Second, the quality-controlled SNPs were further filtered to meet the assumptions of specific analyses, which require unlinked and neutral markers. Using *dartR*, we removed individuals with a more stringent missing data threshold of 20% (i.e. call rate < 0.8), thereby excluding both samples from Ball’s Pyramid and two samples from Roach Island. We then removed monomorphic loci and retained one randomly selected SNP per sequence tag to reduce linkage between markers (removed “secondaries”). Identification and removal of loci out of Hardy-Weinberg equilibrium was performed with all samples combined, using the exact test and an alpha value of 0.01, calculated with the mid-p method. We also checked for loci potentially under selection using BayeScan v.2.1 (Foll & Gaggiotti, 2008), which detects outlier SNPs with significantly elevated pairwise *F*_ST_ values. Following the recommendations of the package, BayeScan was run for 500,000 steps with prior odds of 100, 20 pilot runs and a 10% burn-in. No outliers were detected at the false discovery rate threshold of *q* = 0.05. This (“neutral”) data set comprised 8,012 SNPs from 44 individuals.

### 2.3. Mitogenomic data

We previously generated partial and complete sequences of mitochondrial genomes (“mitogenomes”) from a subset of the same specimens of *P. lata* sampled for SNP generation, encompassing all contemporary and historical populations (Adams et al., 2024; note that the Roach Island samples were all drawn from the 2000 sampling collection). These were combined with mitogenomic sequences from the closely related outgroup species *P. cribrata* and *Panesthia matthewsi*. All data were sourced from GenBank (Supplementary Table S1). We excluded intergenic regions and aligned each gene individually using MUSCLE (Edgar, 2004). We checked for reading frames and premature stop codons in Seqotron v.1.0.1 (Fourment & Holmes, 2016), and removed ambiguously aligned regions. Our final alignment totalled 14,592 bp across 21 samples.

### 2.4. Population structure

We employed four different methods to assess population structure based on the SNP data set. Before proceeding, we checked that the data from Roach Island were not affected by temporal bias in sampling by visualizing intra-population structure with a principal coordinates analysis (PCoA) and calculating pairwise kinship between all individuals (see below for methods). No discernible pattern was found and all samples were retained for analysis.

First, to visually summarize patterns of genetic variation, we conducted a PCoA of individual allele frequencies using *dartR*. In the absence of population genetic assumptions, the PCoA was performed using the quality-controlled data. Before proceeding, we applied a stringent filter for call rate (> 0.95), to account for a high proportion of missing data in the Ball’s Pyramid samples. Remaining analyses were performed using the neutral data set (omitting both individuals from Ball’s Pyramid).

Second, we performed a Bayesian clustering analysis in STRUCTURE v.2.3.4 (Pritchard et al., 2000), automated and parallelized with STRAUTO v.1.0 (Chhatre & Emerson, 2017). We modelled up to six ancestral populations (*K*=1–6), with 10 replicates for each value of *K*, no prior population information, and assuming admixture and independent allele frequencies. Each replicate was run for 500,000 steps with a burn-in of 10%. The optimal *K* value was subsequently selected based on six different metrics calculated in StructureSelector (Li & Liu, 2018). We validated the results using a hierarchical approach, whereby the clusters identified at each step were separated, refiltered and re-analysed independently. The results of replicate runs were summarized and visualized using CLUMPAK (Kopelman et al., 2015) implemented in StructureSelector.

Third, to explore how genetic diversity is partitioned between and within populations, we performed a hierarchical analysis of molecular variance (AMOVA; Excoffier et al., 1992) in GenoDive v.3.0 (Meirmans, 2020) and assessed the significance of results using 10,000 permutations. Populations were defined *a priori* according to sampling locality, which corresponded to genetic partitions delimited by the PCoA and STRUCTURE analyses. Fourth, we estimated genetic divergence between populations by calculating pairwise *F*_ST_ values (Weir & Cockerham, 1984) with the R package *StAMPP* v.1.6.3 (Pembleton et al., 2013). We tested statistical significance with 95% confidence intervals constructed using 10,000 permutations.

To check whether the observed population structure was influenced by the presence of closely related individuals, we calculated pairwise individual kinship coefficients using maximum-likelihood estimation in the R package *SNPRelate* v.1.33.0 (Zheng & Zheng, 2013). We removed one sample from each pair with kinship ≥ 0.125 and re-ran the PCoA, STRUCTURE and *F*_ST_ analyses. Following the removal of relatives, the raw SNP data were filtered with criteria as described above, including a call rate filter of > 0.95 for the analysis including Ball’s Pyramid samples.

### 2.5. Genetic diversity

We investigated genetic health and levels of diversity within the North Bay, Blackburn Island and Roach Island populations. Because the temporally disjunct sampling of Roach Island may have led to biased estimates of inbreeding, we first ran all analyses using the complete data sets, then repeated the procedure using only the 11 Roach Island individuals collected in 2003 (representing the largest single sample). For the latter analyses, the data were re-filtered using methods as above.

We quantified genetic diversity based on the neutral SNP data set by calculating observed and expected heterozygosity, as well as allelic richness, using *hierfstat* v.0.5-11 (Goudet, 2005) and *PopGenReport* v.2.2.2 (Adamack & Gruber, 2014), respectively. Allelic richness was measured with rarefaction to account for uneven sample sizes. To measure inbreeding, we calculated *F*_IS_ values in *hierfstat* and produced 95% confidence intervals in base R. We also estimated kinship coefficients (Loiselle et al., 1995) for each pair of individuals in GenoDive, then averaged these values within and between populations. This method differs from that used in *SNPRelate* by including self-kinship, which is valuable in population diversity estimates (Frankham et al., 2017).

Finally, we estimated effective population size (*N*_e_) from quality-controlled SNPs using the linkage disequilibrium method in Ne-Estimator v.2.1 (Do et al., 2014). Corrections were applied for sampling bias (Waples, 2006) and missing data (Peel et al., 2013). We ran the analysis assuming random mating and report 95% confidence intervals constructed using the jack-knife method, which is more appropriate for large SNP panels (Jones et al., 2016).

### 2.6. Phylogenetic analysis

Phylogenetic analyses were undertaken using the mitogenomic data set. In line with previous work on *Panesthia* cockroaches (see Adams et al., 2024; Beasley-Hall et al., 2021), we divided the data into six partitions: *CO1*, first, second and third codon positions of remaining protein-coding genes, rRNAs, and tRNAs. *CO1* was assigned a separate partition to enable molecular clock calibration (see below), and preliminary analyses confirmed that its segregation did not influence phylogenetic inference. We estimated the maximum-likelihood tree topology in IQTREE2 v.2.2.2 (Minh et al., 2020), using the inbuilt ModelFinder function (Kalyaanamoorthy et al., 2017) to infer the optimal substitution model for each partition based on Bayesian information criterion scores (Supplementary Table S2). Node support was estimated using 10,000 ultrafast bootstrap replicates (UFBoot; Hoang et al., 2018) and the SH-like likelihood-ratio test (SH-aLRT).

We jointly estimated phylogenetic relationships and the evolutionary timescale using a Bayesian framework in BEAST v.1.10.4 (Suchard et al., 2018), with a separate GTR+I+G model applied to each partition. In the absence of any appropriate biogeographic or fossil calibrations, we opted to specify an informative prior for the evolutionary rate of *CO1*. We performed two separate analyses, using the highest and lowest published estimates of molecular rates in insects that diversified during or immediately after the Pleistocene epoch. To obtain a maximum age estimate, in the first analysis we specified a uniform prior of 2.35–3.15 × 10^-2^ substitutions/site/Myr on the rate, based on the dispersal of psyllid plant lice to La Palma island following its formation *ca.* 2 Ma (Percy et al., 2004). To obtain a minimum age estimate, in the second analysis we specified a uniform prior of 1.44–1.73 × 10^-1^ substitutions/site/Myr on the rate, based on the radiation of *Neoconocephalus lyristes* crickets following the opening of the St. Lawrence Seaway *ca.* 11 ka (Ney et al., 2018). In both analyses, we assumed a strict molecular clock, and we compared coalescent (constant size) and birth-death tree priors. Two replicate runs of 2×10^7^ Markov chain Monte Carlo steps were performed for each analysis, drawing samples every 10^4^ steps. Sufficient sampling after convergence was checked using TRACER v.1.7.2 (Rambaut et al., 2018), and maximum-clade-credibility trees were generated in TreeAnnotator with a 10% burn-in.

To check for congruence in the phylogenetic signals between the mitogenomes and nuclear data, we also undertook a phylogenetic analysis of the SNP data set. We filtered the quality-controlled data set for call rate (> 0.95) and secondaries in *dartR*, then concatenated the SNPs to produce an alignment of 1,315 sites. Heterozygous loci were assigned standard ambiguity codes. We inferred the phylogeny using maximum likelihood in IQTREE2 with a GTR+I+G substitution model. Based on results from PCoA analyses, the root was placed between the samples from Ball’s Pyramid and all other taxa.

## 3. Results

### 3.1. Genetic structure

PCoA partitioned genetic variation into four discrete clusters, corresponding to the four contemporary island populations (Figure 2a). The cockroaches from Ball’s Pyramid were especially distant from all remaining populations, and those from Roach Island were somewhat distant from those from Blackburn Island and North Bay.

**Figure 2.**
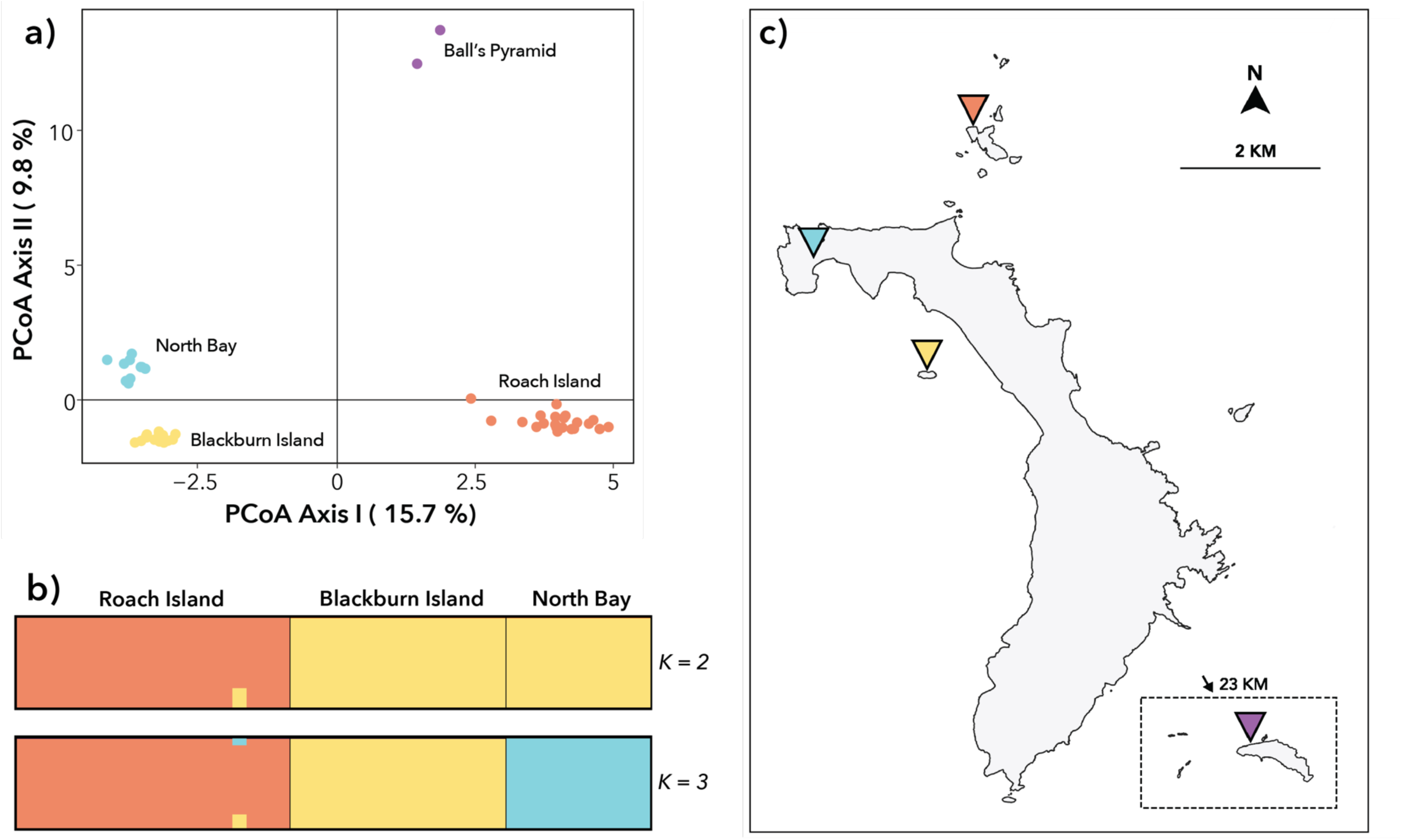
**a)** Principal coordinates analysis (PCoA) plot of 48 samples of *Panesthia lata*, using 1,366 single-nucleotide polymorphisms (SNPs). **b)** STRUCTURE plots for 44 samples based on 8,012 SNPs, when *K* = 2 and *K* = 3 (samples from Ball’s Pyramid were excluded from this analysis). **c)** Map of the Lord Howe Island Group, with labels coloured accordingly to PCoA clusters.

Relationships between the LHI palaeo-island populations (i.e. excluding samples from Ball’s Pyramid) were investigated using sequential STRUCTURE analysis. This suggested that genetic variability was optimally explained by *K* = 2–3 clusters, depending on the estimator, with most supporting *K* = 3 (Supplementary Figure S1). For all values of *K* ≥ 3, there was clear genetic differentiation between the samples from each of the three islands; however when *K* = 2, the samples from North Bay and Blackburn Island collapsed into a single cluster (Figure 2b). Admixture between the populations was negligible irrespective of the *K* value. When samples from each island were analysed separately, no additional population structure was evident.

AMOVA revealed that 54.9% of genetic variance was explained by variation within individuals, while a significant proportion of diversity was explained by variation between islands (24.9%, p < 0.001) and by individuals within islands (20.2%, p < 0.001), indicating appreciable genetic structure. In concordance, pairwise *F*_ST_ values evidenced high, and significant, genetic differentiation between the three populations, with the greatest distance between Roach Island and North Bay (Table 1).

**Table 1.** Genetic differentiation between and within populations of *Panesthia lata* based on 8,012 SNPs. Below diagonal: pairwise *F*_ST_ values (95% confidence interval) (* denotes values significantly different from 0). In bold, on diagonal: average pairwise kinship within populations. Above diagonal: average pairwise kinship between populations.

|  | **Roach Island** | **Blackburn Island** | **North Bay** |
| --- | --- | --- | --- |
| **Roach Island** | **0.199** | -0.105 | -0.116 |
| **Blackburn Island** | 0.239* (0.229–0.250) | **0.202** | -0.011 |
| **North Bay** | 0.279* (0.270–0.289) | 0.225* (0.216–0.234) | **0.328** |

Following the removal of closely related individuals, the data set comprised 12 samples from Blackburn Island, 19 samples from Roach Island and 2 samples from North Bay. PCoA clusters, STRUCTURE clusters and pairwise *F*_ST_ values estimated from this data set were similar to those found using all samples (Supplementary Figure S2; Supplementary Table S3).

### 3.2. Genetic diversity

Average pairwise kinship was high within all populations, with the highest value seen in North Bay and the lowest in Roach Island (Table 1). Average pairwise kinship between all populations was negative, indicating an absence of recent gene flow; however, average kinship was lowest between the populations from Roach Island and North Bay, and highest between those from Blackburn Island and North Bay.

Heterozygosity and allelic richness were highest in the cockroaches from Roach Island and lowest in North Bay, although the differences were relatively modest (Table 2). Further, North Bay was estimated to have a substantially smaller effective population size than Roach Island (Table 2), albeit with broad confidence intervals. The effective population size of Blackburn Island could not be estimated, likely due to insufficient sample size (Do et al., 2014). Estimates of *F*_IS_ were high and significant in all populations, indicating considerable inbreeding (Table 2). In contrast to other metrics of diversity, the estimated *F*_IS_ was highest in Roach Island and lowest in Blackburn Island. Results were similar for all metrics when analyses were repeated using only the 11 Roach Island specimens collected in 2003 (Supplementary Table S4).

**Table 2.**
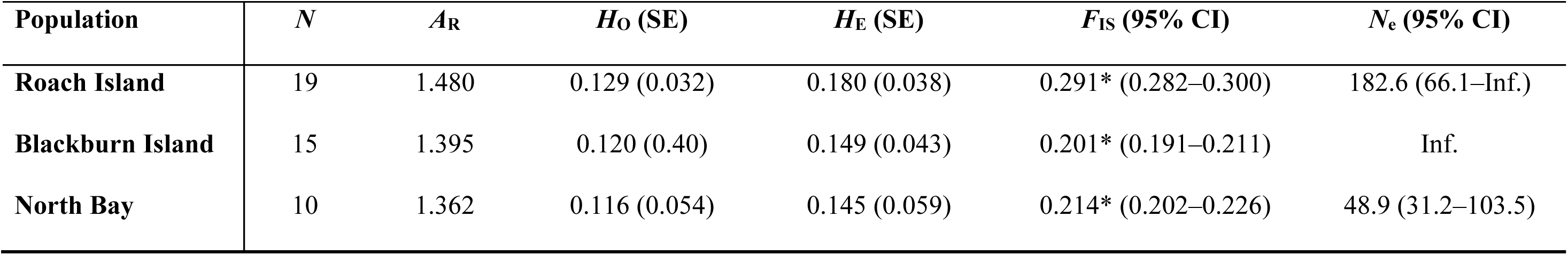
Summary of genetic diversity statistics across three populations of *Panesthia lata*. *N* = sample size, *A*_R_ = mean rarefied allelic richness, *H*_O_ = mean observed heterozygosity, SE = standard error, *H*_O_ = mean expected heterozygosity, *F*_IS_ = inbreeding coefficient (* denotes value significantly different to 0), 95% CI = 95% lower and upper confidence interval, *N*_e_ = effective population size. All values except *N*_e_ based on 8,012 SNPs, *N*_e_ values based on 10,370 SNPs.

### 3.3. Phylogenetic analyses and evolutionary timescale

Maximum-likelihood and Bayesian analyses of complete mitogenomes both placed the Ball’s Pyramid population as the sister group to the remaining lineages with high support (UFBoot, SH-aLRT, PP = 1; Figures 3, 4). However, relationships between and within the other populations were inconsistent. In the maximum-likelihood phylogeny, samples from Roach Island formed a paraphyletic grade with respect to those from Blackburn Island, North Bay and historical LHI, albeit with equivocal node support (Figure 3). Individuals from Blackburn Island and North Bay formed two reciprocally monophyletic clades, although both were nested among historical LHI samples, once more with relatively poor node support.

**Figure 3.**
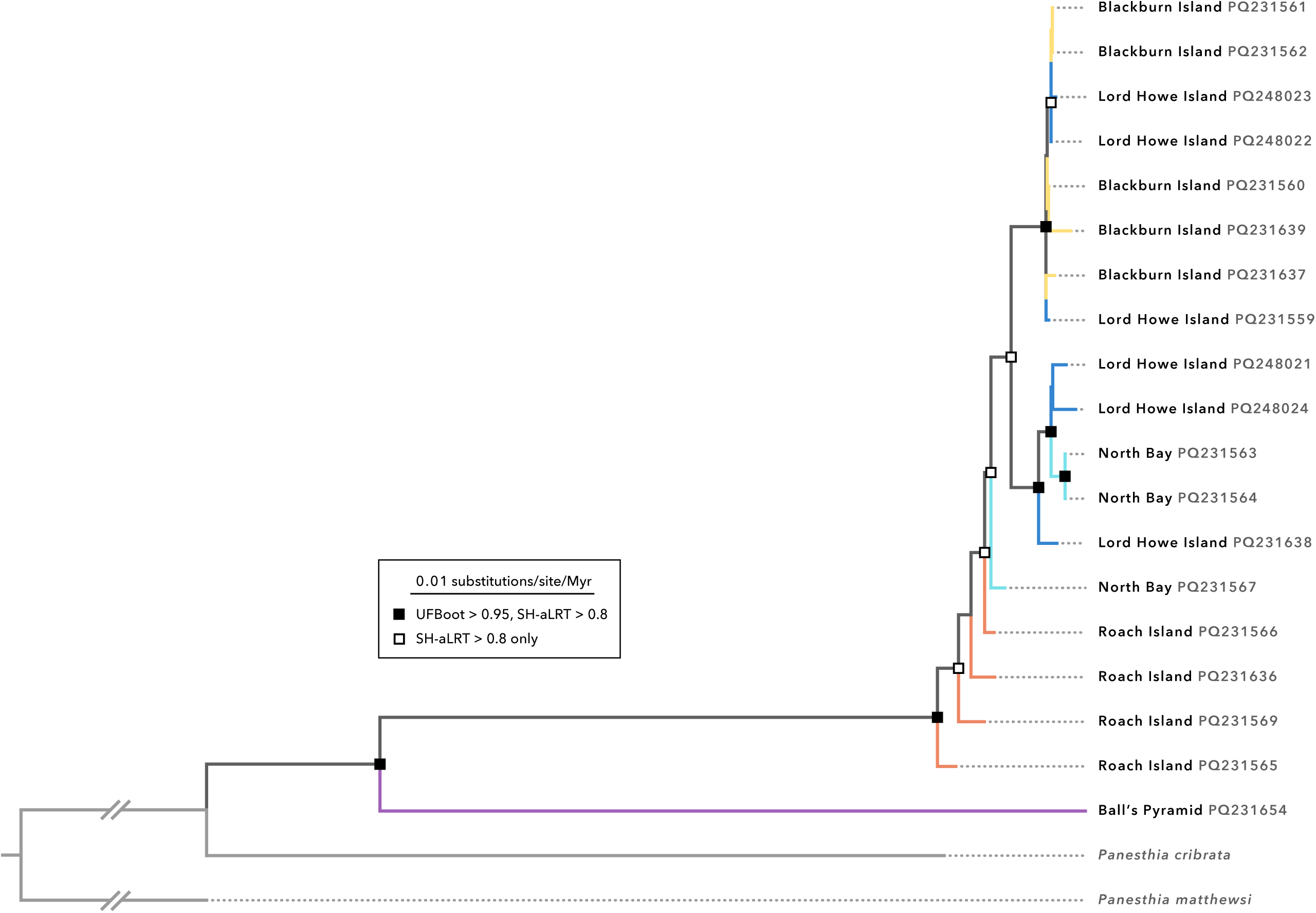
Maximum-likelihood phylogeny of *Panesthia lata* inferred from complete mitochondrial genomes in IQTREE2. UFBoot: ultrafast bootstrap; SH-aLRT: SH-like likelihood-ratio test. Numbers next to tip labels indicate GenBank accession numbers. Specimens labelled with “Lord Howe Island” are historical samples collected prior to near-extirpation by rodents.

**Figure 4.**
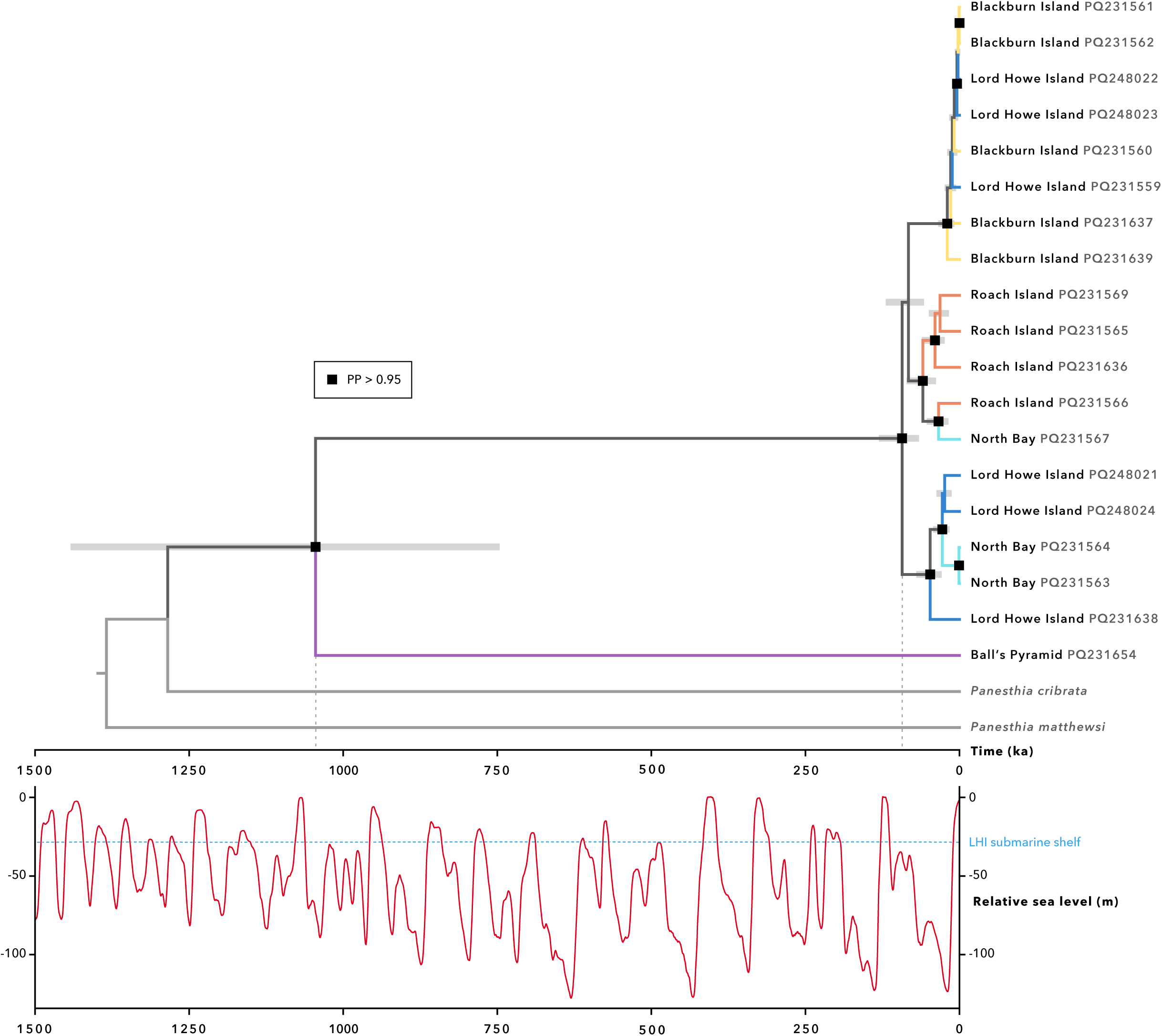
Dated phylogeny of *Panesthia lata* inferred from complete mitochondrial genomes in BEAST. The figure depicts our maximum age estimate, which was obtained by specifying an informative prior on the substitution rate of *CO1* (Percy et al., 2004). PP: posterior probability. Numbers on labels indicate GenBank accession numbers. **Inset:** Relative sea level over the last 1.5 Myr, adapted from Bintanja and van de Wal (2008). Dashed line indicates the median contemporary depth of the Lord Howe palaeo-island submarine shelf (Kennedy et al., 2002). The divergence of the Lord Howe, Blackburn and Roach Island samples initiated during the most recent glacial period.

Bayesian analysis yielded a contrasting topology, inferring a closer relationship between cockroaches from Blackburn Island and Roach Island (Figure 4). With the exception of the individuals from Ball’s Pyramid, none of the populations were recovered as monophyletic. Individuals from North Bay and Blackburn Island once more grouped with historical LHI specimens, with variable node support. Curiously, the Roach Island samples formed a well-supported clade that was paraphyletic with respect to a single individual from North Bay.Using a low and high estimate of the substitution rate for CO1, we obtained estimates of divergence times that we consider to be maximum and minimum values, respectively. Using the lower estimate of the substitution rate, the cockroaches from Ball’s Pyramid were inferred to have diverged from the remaining populations *ca.* 1.04 Ma (95% highest posterior density interval [HPD] 0.784–1.44 Ma), while the crown age of the LHI palaeo-island clade was found to be *ca.* 93.1 ka (95% HPD 67.4–129 ka; Figure 4). Using the higher estimate for the substitution rate yielded shallower and narrower date ranges (Supplementary Figure S3). The divergence of Ball’s Pyramid samples was inferred to have occurred *ca.* 185 ka (95% HPD 133–242 ka) and the crown age of the LHI palaeo-island was estimated at *ca.* 16.5 ka (95% HPD 12.0–21.8 ka). These estimates were derived using a coalescent tree prior, and values were nearly identical under a birth-death tree prior.

Finally, maximum-likelihood analysis of nuclear SNPs resolved all populations as reciprocally monophyletic (Supplementary Figure S4). The samples from Roach Island formed a well-supported sister group to those from Blackburn Island and North Bay (UFBoot, SH-aLRT = 1).

## 4. Discussion

### 4.1. Phylogeography and sea level

The pattern of genetic structure in *P. lata* is consistent with the geographic arrangement of palaeo-islands, and suggests that land-bridge connectivity allowed migration over the LHI palaeo-island. Across all analyses of mitochondrial and nuclear data, the population on Ball’s Pyramid was especially divergent from the remaining samples, reflecting its permanent geographic isolation from the LHI palaeo-island (Gilmore et al., 2020; McDougall et al., 1981). While we acknowledge a higher proportion of missing data for the Ball’s Pyramid specimens, our results were robust to data filtering, indicating that the population’s distinctiveness was not artefactual. In concordance, our divergence date estimates range from 185 ka to 1.04 Ma, suggesting that the population has been isolated over multiple sea-level cycles. Here we provide the first genetic evidence of the evolutionary distinctiveness of the Ball’s Pyramid fauna (Flemons et al., 2018; Lilemets & Wilson, 2002), as well as its connection to the island’s geological and oceanographic context. It remains unclear how *P. lata* reached the Pyramid, but a plausible explanation involves rafting, potentially when the island’s subaerial extent was larger during a glacial period.

The weaker differentiation among the remaining populations points to more recent gene flow, presumably via overland migration during sea-level lowstands. Our time-calibrated phylogenies indicate that the LHI palaeo-island populations began to diverge 16.6 to 93.1 ka. Due to the non-monophyly of mitochondrial lineages and the recency of isolation, it is challenging to achieve precise molecular date estimates, and these are likely to be overestimates of the true age of geographic separation (Edwards & Beerli, 2000; Giarla & Esselstyn, 2015). However, this approximate timescale places the genetic coalescence during the previous glacial period (100–10 ka; Spratt & Lisiecki, 2016), which is consistent with dispersal across the most recent land bridges. Importantly, this result is highly inconsistent with the species pump hypothesis (Heaney, 1985) or the hypothesis that genetic structure may not reflect physical connectivity (e.g., Cros et al., 2020). Instead, we demonstrate that the LHIG is another archipelago for which Pleistocene land bridges facilitated gene flow (Barker et al., 2012; Fattorini, 2010; Fernández-Palacios et al., 2015; Heaney et al., 2005; Papadopoulou & Knowles, 2017).

The Roach Island population was relatively distant from the North Bay and Blackburn Island populations in nuclear SNP analyses, as well as in the maximum-likelihood mitochondrial phylogeny, which included historical LHI samples. The channel isolating Roach Island is deeper and broader than that between LHI and Blackburn Island, hence this result further implicates the recent sea-level inundation as a key driver of genetic structure in *P. lata*. However, considering the uncertainty in our molecular date estimates, we cannot exclude the possibility that divergence initiated prior to end-Pleistocene sea-level rise (*ca.* 10 ka; Spratt & Lisiecki, 2016). In which case, genetic structure could also have been shaped by other barriers to gene flow, such as the heterogeneous topography and/or biotic landscape of LHI (Hyman et al., 2023; Major et al., 2021). The relative influences of sea level versus topography are difficult to tease apart without granular geographic sampling across LHI itself.

Our estimated evolutionary timescale stands in sharp contrast to previous estimates of the age of the Blackburn and Roach Island populations (*ca.* 3 Ma in Lo et al., 2016; *ca.* 330 ka in Adams et al., 2024). Several additional lines of evidence support the shallow timescale inferred presently. First, the dates inferred by Lo et al. (2016) and Adams et al. (2024) were based on deep fossil calibrations and a substitution rate calibrated on a more ancient timescale, respectively (and with only two samples of *P. lata* included in the phylogeny of Lo et al., 2016). It is now well established that temporally distant calibrations can lead to substantial overestimation of population-level coalescence times (Grant, 2015; Ho et al., 2011). Second, the non-monophyly in our mitogenome analyses, even when discounting the historical LHI specimens, is more likely to reflect incomplete lineage sorting following recent rather than ancient isolation (e.g., Brüniche-Olsen et al., 2021; Li et al., 2020). Third, while genetic studies across the LHIG are scarce, a previous phylogenetic analysis found minimal differentiation between individuals of the skink *Oligosoma lichenigerum* from Blackburn Island and LHI based on several nuclear and mitochondrial markers (Chapple et al., 2009). Taken together, these and our results support the inference that recent glacial land bridges have been permeable to gene flow. In future, molecular dating could be refined by additional evidence, such as a species-specific estimates of the evolutionary rate.

Overall, notwithstanding the uncertainty in the molecular date estimates, the broad patterns of genetic structure in *P. lata* remain consistent with gene flow across the LHI palaeo-island at some point in the Pleistocene epoch. This contrasts with suggestions that there has been sustained isolation and *in situ* speciation on the islets, producing highly divergent ecological communities (Cassis et al., 2003; Lillemets & Wilson, 2002). Indeed, since *P. lata* is an ecologically specialized and non-vagile species, and unlikely to readily disperse across land bridges, we expect that the present findings can be generalized to other organisms.

### 4.2. Genetic structure and relict populations

Our nuclear SNP analyses reveal four contemporary genetic clusters within *P. lata*, corresponding to the four islands that compose its archipelagic range. Most significantly, the genetic distinctiveness of the North Bay population confirms its status as a relic of the former diversity on LHI, as opposed to a recent transplant from another islet. By accounting for closely related individuals in PCoA and STRUCTURE analysis, we ensured that this result was not an artefact of high kinship (e.g., Ewart et al., 2021). That the population survived more than a century of rodent activity is remarkable, even if its size and spatial extent are highly limited. One possible explanation is that predation was limited by rodent control in an adjacent *Howea* palm plantation, where baiting and trapping have been underway since the 1940s (Harper et al., 2020; D. Hiscox, pers. comm.). Although we were unable to generate genetic data for the second LHI population, its geographic distance from North Bay suggests that it also represents a separate relict population. Moreover, since our surveys were non-exhaustive, the persistence of *P. lata* at this site raises the possibility of further relict populations being discovered in future, as the species recovers following predator release.

Interestingly, despite their clear partitioning based on nuclear SNPs, the cockroaches from Roach Island, Blackburn Island and North Bay all emerged as non-monophyletic in mitogenomic analyses. Considering the recency of divergence, this discordance could be attributable to introgression, selection upon mitochondrial variants or incomplete lineage sorting. It is not unusual for genomic SNP panels to detect fine-scale population structure not yet manifest in the genealogies of individual loci (e.g., McCartney-Melstad et al., 2018; Sturm et al., 2020; Younger et al., 2017). However, a significant factor in the mitochondrial non-monophyly was the inclusion of historical LHI samples, which were not genotyped for nuclear SNPs. One explanation is that the pre-rodent population on LHI comprised a wider diversity of haplotypes, some with close affinity to the other islets, which were subsequently lost as the population declined. Alternatively, while the historical specimens were all labelled as originating from “Lord Howe Island”, some may have been collected from Blackburn Island, which is near to the human settlement. Individuals from the two populations cannot be reliably separated by morphology, thus further sampling of nuclear markers will be needed to confirm the provenance of the ancient specimens.

### 4.3. Genetic diversity and conservation implications

The discovery that *P. lata* has survived on LHI is a significant advancement in the species’ conservation, yet it does not preclude the need for ongoing management. Genetic diversity and effective population size were low in all three sampled populations, consistent with their small geographic extent and insularity. Although the small sample sizes and broad confidence intervals limit interpretation, our estimates of inbreeding far exceed the levels at which other species suffer large reductions in fitness and survivorship, including insects (Frankham et al., 2017; Grueber et al., 2008; Huisman et al., 2016; Müller & Juškauskas, 2018; Ralls et al., 2018). Our results therefore evince the need for ongoing monitoring of the species, to enable prompt intervention if populations decline.

Our study has also provided the first comparison of genetic diversity across different islands of the LHIG. Across most metrics, genetic health and diversity were highest among the cockroaches in Roach Island and lowest in North Bay. This pattern reflects the extent and quality of habitat, whereby Roach Island (the largest islet after Ball’s Pyramid) has been mostly undisturbed by human activity, while large sections of Blackburn Island have been deforested and LHI has been severely impacted by invasive rodents (DECC, 2007; Priddel & Wheeler, 2014). It is therefore curious that *F*_IS_ was highest in Roach Island, which could indicate a recent uptick in inbreeding. While the genetic health of the Roach Island population warrants further study, the high kinship and excess of close relatives within the North Bay population are of particular concern, establishing the relict LHI population as a conservation priority.

Prior to the discovery of the relict population, several authors had proposed a reintroduction of *P. lata* to LHI (Carlile et al., 2018; Hutton et al., 2007). Although reintroduction is now unnecessary, the vulnerability of the population raises the possibility of genetic rescue: the transplanting of propagules from elsewhere to counteract genetic erosion (Tallmon et al., 2004; Whiteley et al., 2015). Typically, due to the risk of outbreeding depression, genetic rescues are undertaken between populations that have been isolated for under 500 years (Frankham et al., 2011). However, this threshold is likely to be conservative (Ralls et al., 2018), and studies evidence long-term fitness increases even when populations diverged at a comparable or earlier age than *P. lata* (Aitken & Whitlock, 2013; Harrisson et al., 2016; Kronenberger et al., 2017; Pekkala et al., 2012; Weeks et al., 2017). Nonetheless, further data will be needed to assess the risk of outbreeding depression, including captive-crossing or comparative genomic studies.

Finally, we confirm the presence of *P. lata* on Ball’s Pyramid, which is not currently recognized in the species’ management plan due to the islet’s inaccessibility for survey. Based on its significant divergence in nuclear allele frequencies and evolutionary distinctiveness (inferred from phylogenetic analysis), the population could qualify as an evolutionarily significant unit (ESU; an independently evolving, divergent unit of genetic variance; Moritz, 1994; Zink, 2004). ESU status could grant legal protection and encourage focused management of the unique habitat (reviewed by Funk et al., 2012). While we were unable to measure genetic diversity on Ball’s Pyramid, ecological observations suggest that *P. lata* has a limited presence on the island and may therefore be vulnerable to local extinction (N. Carlile, pers. obs.). Overall, our findings reveal that the natural and anthropogenic fragmentation of the LHIG has significantly impacted genetic diversity in *P. lata*, and have clarified priorities for the species’ future management.

## 5. Conclusions

The impact of Pleistocene sea-level fluctuation remains a fertile and debated question in island biogeography. This study represents the first detailed, species-wide genetic analysis across multiple islands of the LHIG, broadening the field with the addition of a novel model system. Our results offer little support for the hypothesis that sea-level fluctuation could have acted as a species pump on the LHIG, or that genetic structure was unaffected by sea-level change. Instead, we demonstrate that the LHIG is another archipelago that conforms to the traditional model of palaeo-island biogeography, whereby genetic structure within *P. lata* reflects the proximity and historic fusion of some islets into a larger landmass, allowing for end-Pleistocene gene flow.

From a conservation standpoint, our discovery of a highly evolutionary distinct lineage on Ball’s Pyramid reiterates the need to conserve the islet’s unique habitat and fauna, which appear to have been long isolated from LHI. Moreover, the relatively shallow divergence of the Lord Howe, Blackburn and Roach Island populations, alongside the high inbreeding in these populations, provides evidence that genetic rescue between these islands may be an effective conservation strategy for *P. lata*. Though further work will be needed to confirm whether these findings can be generalized, we suggest that these results could also inform the management of other species with similar distributions across the LHIG.

## Acknowledgements

This project was supported by the Australia Pacific Science Foundation (grant APSF22029), and samples were collected under permits from NSW National Parks and Wildlife Services (SL102663) and the Lord Howe Island Board (05/22). The authors acknowledge the Sydney Informatics Hub and the University of Sydney’s high-performance computing cluster, Artemis, for computational resources that contributed to results in the present paper. We thank Chris Reid, Derek Smith, Jude Philp and Matthew Hahn for access to historical specimens, as well as Hank Bower, Jack Schick, Dean Hiscox, Justin Gilligan, Caitlin Woods and the Lord Howe Island Board for support and assistance with fieldwork. Justin Gilligan also provided the photograph of *P. lata*, for which we are very grateful. Finally, we thank Toby Kovacs, Ros Gloag and Carolyn Hogg for valuable comments on an earlier draft of the manuscript.

## Data availability statement

Raw single-nucleotide polymorphism data, phylogenetic trees and tables are to be deposited in Dryad.

## Data accessibility

SNP data used in this study are to be uploaded to Dryad, and mitogenomes were sourced from GenBank.

## Author contributions

Maxim W.D. Adams: original conception, sampling, data analysis, writing original draft, reviewing; Kyle M. Ewart: data analysis, reviewing; Nicholas Carlile: sampling, reviewing, supervision; Harley A. Rose: sampling, reviewing, supervision; James A. Walker: reviewing, supervision; Ian Hutton: sampling, reviewing; Simon Y.W. Ho: reviewing, supervision; Nathan Lo: original conception, sampling, reviewing, supervision.

## References

Adamack, A. T., & Gruber, B. (2014). PopGenReport: simplifying basic population genetic analyses in R. Methods in Ecology and Evolution, 5(4), 384–387.

Aitken, S. N., & Whitlock, M. C. (2013). Assisted gene flow to facilitate local adaptation to climate change. Annual Review of Ecology, Evolution, and Systematics, 44, 367–388.

Ali, J.R., & Aitchison, J.C. (2014). Exploring the combined role of eustasy and oceanic island thermal subsidence in shaping biodiversity on the Galápagos. Journal of Biogeography, 41, 1227–1241. 10.1111/jbi.12313

Anderson, A. (2003). Investigating early settlement on Lord Howe Island. Australian Archaeology, 57(1), 98–102. 10.1080/03122417.2003.11681767

Ávila, S.P., Melo, C., Berning, B., Sá, N., Quartau, R., Rijsdijk, K.F., Ramalho, R.S., Cordeiro, R., De Sá, N.C., Pimentel, A., Baptista, L., Medeiros, A., Gil, A., & Johnson, M.E. (2019). Towards a ‘Sea-Level Sensitive’ dynamic model: impact of island ontogeny and glacio-eustasy on global patterns of marine island biogeography. Biological Reviews, 94(3), 1116–1142. 10.1111/brv.12492

Barker, B. S., Rodríguez-Robles, J. A., Aran, V. S., Montoya, A., Waide, R. B., & Cook, J. A. (2012). Sea level, topography and island diversity: phylogeography of the Puerto Rican Red-eyed Coquí, *Eleutherodactylus antillensis*. Molecular Ecology, 21(24), 6033–6052. 10.1111/mec.12020

Bintanja, R., & van de Wal, R. S. W. (2008). North American ice-sheet dynamics and the onset of 100,000-year glacial cycles. Nature, 454(7206), 869–872. 10.1038/nature07158

Brown, R.M., Siler, C.D., Oliveros, C.H., Esselstyn, J.A., Diesmos, A.C., Hosner, P.A., Linkem, C.W., Barley, A.J., Oaks, J.R., Sanguila, M.B., & Welton, L.J. (2013). Evolutionary processes of diversification in a model island archipelago. Annual Review of Ecology, Evolution, and Systematics, 44(1), 411–435. 10.1146/annurev-ecolsys-110411-160323

Brüniche-Olsen, A., Bickham, J. W., Godard-Codding, C. A., Brykov, V. A., Kellner, K. F., Urban, J., & DeWoody, J. A. (2021). Influence of Holocene habitat availability on Pacific gray whale (*Eschrichtius robustus*) population dynamics as inferred from whole mitochondrial genome sequences and environmental niche modeling. Journal of Mammalogy, 102(4), 986–999.

Carlile, N., & Priddel, D. (2013a). Seabird islands No. 257: Tenth of June Island, Lord Howe Group, New South Wales. Corella, 37(4), 86–87.

Carlile, N., & Priddel, D. (2013b). Seabird islands No. 259: Sugarloaf Island, Lord Howe Group, New South Wales. Corella, 37(4), 90–91.

Carlile, N., & Priddel, D. (2013c). Seabird islands No. 260: Soldiers Cap, Lord Howe Group, New South Wales. Corella, 37(4), 92–93.

Carlile, N., & Priddel, D. (2013d). Seabird islands No. 261: Mutton Bird Island, Lord Howe Group, New South Wales. Corella, 37(4), 94–96.

Carlile, N., & Priddel, D. (2013e). Seabird islands No. 262: Blackburn Island, Lord Howe Group, New South Wales. Corella, 37(4), 97–99.

Carlile, N., Priddel, D., & Bower, H. (2013). Seabird islands No. 256: Roach Island, Lord Howe Group, New South Wales. Corella, 37(4), 82–85.

Carlile, N., Priddel, D., & O’Dwyer, T. (2018). Preliminary surveys of the endangered Lord Howe Island cockroach *Panesthia lata* (Blattodea: Blaberidae) on two islands within the Lord Howe Group, Australia. Austral Entomology, 57(2), 207–213. 10.1111/aen.12281

Cassis, G., Meades, L., Reid, C., Harris, R., Carter, G., & Jefferys, E. (2003). Lord Howe Island: Terrestrial Invertebrate Biodiversity and Conservation. A report prepared by the Australian Museum Centre for Biodiversity Conservation and Research for NSW National Parks.

Chapple, D. G., Ritchie, P. A., & Daugherty, C. H. (2009). Origin, diversification, and systematics of the New Zealand skink fauna (Reptilia: Scincidae). Molecular Phylogenetics and Evolution, 52(2), 470–487.

Chhatre, V. E., & Emerson, K. J. (2017). StrAuto: automation and parallelization of STRUCTURE analysis. BMC Bioinformatics, 18(1), 192.

Cros, E., Chattopadhyay, B., Garg, K. M., Ng, N. S. R., Tomassi, S., Benedick, S., Edwards, D. P., & Rheindt, F. E. (2020). Quaternary land bridges have not been universal conduits of gene flow. Molecular Ecology, 29(14), 2692–2706. 10.1111/mec.15509

Cros, E., & Rheindt, F. E. (2017). Massive bioacoustic analysis suggests introgression across Pleistocene land bridges in *Mixornis* tit-babblers. Journal of Ornithology, 158, 407–419.

Cruz, V. M. V., Kilian, A., & Dierig, D. A. (2013). Development of DArT marker platforms and genetic diversity assessment of the US collection of the new oilseed crop lesquerella and related species. PLOS ONE, 8(5), e64062.

De Meeûs, T. (2018). Revisiting *F*_IS_, *F*_ST_, Wahlund effects, and null alleles. Journal of Heredity, 109(4), 446–456.

DECC (2007). Lord Howe Island biodiversity management plan. Department of Environment and Climate Change NSW.

Do, C., Waples, R. S., Peel, D., Macbeth, G. M., Tillett, B. J., & Ovenden, J. R. (2014). NeEstimator v2: re-implementation of software for the estimation of contemporary effective population size (Ne) from genetic data. Molecular Ecology Resources, 14(1), 209–214. 10.1111/1755-0998.12157

Edgar, R. C. (2004). MUSCLE: multiple sequence alignment with high accuracy and high throughput. Nucleic Acids Research, 32(5), 1792–1797.

Edwards, S., & Beerli, P. (2000). Perspective: gene divergence, population divergence, and the variance in coalescence time in phylogeographic studies. Evolution, 54(6), 1839–1854.

Ewart, K. M., Ho, S. Y. W., Chowdhury, A.-A., Jaya, F. R., Kinjo, Y., Bennett, J., Bourguignon, T., Rose, H. A., & Lo, N. (2024). Pervasive relaxed selection in termite genomes. Proceedings of the Royal Society B, 291(2023), 20232439.

Ewart, K. M., Johnson, R. N., Joseph, L., Ogden, R., Frankham, G. J., & Lo, N. (2021). Phylogeography of the iconic Australian pink cockatoo, *Lophochroa leadbeateri*. Biological Journal of the Linnean Society, 132(3), 704–723.

Excoffier, L., Smouse, P. E., & Quattro, J. (1992). Analysis of molecular variance inferred from metric distances among DNA haplotypes: application to human mitochondrial DNA restriction data. Genetics, 131(2), 479–491.

Fattorini, S. (2010). Influence of Recent geography and palaeogeography on the structure of reptile communities in a land-bridge archipelago. Journal of Herpetology, 44(2), 242–252, 211. 10.1670/09-046.1

Fernández-Palacios, J.M., Rijsdijk, K.F., Norder, S.J., Otto, R., de Nascimento, L., Fernández-Lugo, S., Tjørve, E., & Whittaker, R.J. (2016). Towards a glacial-sensitive model of island biogeography. Global Ecology and Biogeography, 25, 817–830. 10.1111/geb.12320

Foll, M., & Gaggiotti, O. (2008). A genome-scan method to identify selected loci appropriate for both dominant and codominant markers: a Bayesian perspective. Genetics, 180(2), 977–993. 10.1534/genetics.108.092221

Fourment, M., & Holmes, E. C. (2016). Seqotron: a user-friendly sequence editor for Mac OS X. BMC Research Notes, 9, 106.

Frankham, R., Ballou, J. D., Eldridge, M. D., Lacy, R. C., Ralls, K., Dudash, M. R., & Fenster, C. B. (2011). Predicting the probability of outbreeding depression. Conservation Biology, 25(3), 465–475.

Frankham, R., Ballou, J. D., Ralls, K., Eldridge, M., Dudash, M. R., Fenster, C. B., Lacy, R. C., & Sunnucks, P. (2017). Genetic management of fragmented animal and plant populations. Oxford University Press.

Funk, W. C., McKay, J. K., Hohenlohe, P. A., & Allendorf, F. W. (2012). Harnessing genomics for delineating conservation units. Trends in Ecology & Evolution, 27(9), 489–496.

Garg, K. M., Chattopadhyay, B., Wilton, P. R., Malia Prawiradilaga, D., & Rheindt, F. E. (2018). Pleistocene land bridges act as semipermeable agents of avian gene flow in Wallacea. Molecular Phylogenetics and Evolution, 125, 196–203. 10.1016/j.ympev.2018.03.032

Giarla, T. C., & Esselstyn, J. A. (2015). The challenges of resolving a rapid, recent radiation: empirical and simulated phylogenomics of Philippine shrews. Systematic Biology, 64(5), 727–740.

Gilmore, P., Bull, K., Matchan, E., Phillips, D., Hutton, I., Coenraads, R., Roots, W., & Higgins, R. (2020). New Ar-Ar geochronology age constraints for volcanism on Lord Howe Island. *Geological Survey of New South Wales Report*, GS2020/0503, 1–23.

Goudet, J. (2005). HIERFSTAT, a package for R to compute and test hierarchical *F*-statistics. Molecular Ecology Notes, 5(1), 184–186.

Grant, W. S. (2015). Problems and cautions with sequence mismatch analysis and Bayesian skyline plots to infer historical demography. Journal of Heredity, 106(4), 333–346.

Green, P. (1993). Notes relating to the floras of Norfolk & Lord Howe Islands, IV. Kew Bulletin, 48(2), 307–325.

Gruber, B., Unmack, P. J., Berry, O. F., & Georges, A. (2018). DARTR: An R package to facilitate analysis of SNP data generated from reduced representation genome sequencing. Molecular Ecology Resources, 18(3), 691–699.

Grueber, C. E., Wallis, G. P., & Jamieson, I. G. (2008). Heterozygosity–fitness correlations and their relevance to studies on inbreeding depression in threatened species. Molecular Ecology, 17(18), 3978–3984. 10.1111/j.1365-294X.2008.03910.x

Harper, G. A., Pahor, S., & Birch, D. (2020). The Lord Howe Island rodent eradication: lessons learnt from an inhabited island. Proceedings of the Vertebrate Pest Conference.

Harrisson, K. A., Pavlova, A., Gonçalves da Silva, A., Rose, R., Bull, J. K., Lancaster, M. L., Murray, N., Quin, B., Menkhorst, P., Magrath, M. J. L., & Sunnucks, P. (2016). Scope for genetic rescue of an endangered subspecies though re-establishing natural gene flow with another subspecies. Molecular Ecology, 25(6), 1242–1258. 10.1111/mec.13547

Heaney, L. R. (1985). Zoogeographic evidence for middle and late Pleistocene land bridges to the Philippine Islands. Modern Quaternary Research in Southeast Asia, 9, 127–144.

Heaney, L. R., Walsh Jr, J. S., & Townsend Peterson, A. (2005). The roles of geological history and colonization abilities in genetic differentiation between mammalian populations in the Philippine archipelago. Journal of Biogeography, 32(2), 229–247. 10.1111/j.1365-2699.2004.01120.x

Hedrick, P. W., & Garcia-Dorado, A. (2016). Understanding inbreeding depression, purging, and genetic rescue. Trends in Ecology & Evolution, 31(12), 940–952.

Ho, S. Y. W., Lanfear, R., Bromham, L., Phillips, M. J., Soubrier, J., Rodrigo, A. G., & Cooper, A. (2011). Time-dependent rates of molecular evolution. Molecular Ecology, 20(15), 3087–3101.

Hoang, D. T., Chernomor, O., Von Haeseler, A., Minh, B. Q., & Vinh, L. S. (2018). UFBoot2: improving the ultrafast bootstrap approximation. Molecular Biology and Evolution, 35(2), 518–522.

Huisman, J., Kruuk, L. E., Ellis, P. A., Clutton-Brock, T., & Pemberton, J. M. (2016). Inbreeding depression across the lifespan in a wild mammal population. Proceedings of the National Academy of Sciences of the U.S.A., 113(13), 3585–3590.

Hutton, I., Parkes, J. P., & Sinclair, A. R. (2007). Reassembling island ecosystems: the case of Lord Howe Island. Animal Conservation, 10(1), 22–29.

Hyman, I. T., Caiza, J., & Köhler, F. (2023). Dissecting an island radiation: systematic revision of endemic land snails on Lord Howe Island (Gastropoda: Stylommatophora: Microcystidae). Zoological Journal of the Linnean Society, 197(1), 20–75.

Jin, M., Zwick, A., Ślipiński, A., de Keyzer, R., & Pang, H. (2020). Museomics reveals extensive cryptic diversity of Australian prionine longhorn beetles with implications for their classification and conservation. Systematic Entomology, 45(4), 745–770. 10.1111/syen.12424

Jones, A., Ovenden, J., & Wang, Y. (2016). Improved confidence intervals for the linkage disequilibrium method for estimating effective population size. Heredity, 117(4), 217–223.

Kalyaanamoorthy, S., Minh, B. Q., Wong, T. K., Von Haeseler, A., & Jermiin, L. S. (2017). ModelFinder: fast model selection for accurate phylogenetic estimates. Nature Methods, 14(6), 587–589.

Kardos, M., Zhang, Y., Parsons, K. M., Kang, H., Xu, X., Liu, X., Matkin, C. O., Zhang, P., Ward, E. J., & Hanson, M. B. (2023). Inbreeding depression explains killer whale population dynamics. Nature Ecology & Evolution, 7, 657–686.

Kennedy, E. S., Grueber, C. E., Duncan, R. P., & Jamieson, I. G. (2014). Severe inbreeding depression and no evidence of purging in an extremely inbred wild species—the Chatham Island black robin. Evolution, 68(4), 987–995.

Kilian, A., Wenzl, P., Huttner, E., Carling, J., Xia, L., Blois, H., Caig, V., Heller-Uszynska, K., Jaccoud, D., Hopper, C., Aschenbrenner-Kilian, M., Evers, M., Peng, K., Cayla, C., Hok, P., & Uszynski, G. (2012). Diversity arrays technology: a generic genome profiling technology on open platforms. Methods in Molecular Biology, 888, 67–89. 10.1007/978-1-61779-870-2_5

Kopelman, N. M., Mayzel, J., Jakobsson, M., Rosenberg, N. A., & Mayrose, I. (2015). Clumpak: a program for identifying clustering modes and packaging population structure inferences across K. Molecular Ecology Resources, 15(5), 1179–1191.

Kronenberger, J. A., Funk, W. C., Smith, J. W., Fitzpatrick, S. W., Angeloni, L. M., Broder, E. D., & Ruell, E. W. (2017). Testing the demographic effects of divergent immigrants on small populations of Trinidadian guppies. Animal Conservation, 20(1), 3–11. 10.1111/acv.12286

Leonard, J.A., den Tex, R.-J., Hawkins, M.T.R., Muñoz-Fuentes, V., Thorington, R., & Maldonado, J.E. (2015). Phylogeography of vertebrates on the Sunda Shelf: a multi-species comparison. Journal of Biogeography, 42, 871–879. f10.1111/jbi.12465

Li, F., & Li, S. (2018). Paleocene–Eocene and Plio–Pleistocene sea-level changes as “species pumps” in Southeast Asia: evidence from *Althepus* spiders. Molecular Phylogenetics and Evolution, 127, 545–555. 10.1016/j.ympev.2018.05.014

Li, K., Ren, X., Song, X., Li, X., Zhou, Y., Harlev, E., Sun, D., & Nevo, E. (2020). Incipient sympatric speciation in wild barley caused by geological-edaphic divergence. Life Science Alliance, 3(12), e202000827.

Li, Y. L., & Liu, J. X. (2018). StructureSelector: A web-based software to select and visualize the optimal number of clusters using multiple methods. Molecular Ecology Resources, 18(1), 176–177.

Lillemets, B., & Wilson, G. D. (2002). Armadillidae (Crustacea: Isopoda) from Lord Howe Island: new taxa and biogeography. Records of the Australian Museum, 54(1), 71–98.

Lo, N., Tong, K. J., Rose, H. A., Ho, S. Y., Beninati, T., Low, D. L., Matsumoto, T., & Maekawa, K. (2016). Multiple evolutionary origins of Australian soil-burrowing cockroaches driven by climate change in the Neogene. Proceedings of the Royal Society B, 283(1825), 20152869.

Loiselle, B. A., Sork, V. L., Nason, J., & Graham, C. (1995). Spatial genetic structure of a tropical understory shrub, *Psychotria officinalis* (Rubiaceae). American Journal of Botany, 82(11), 1420–1425.

Lord Howe Island Board (2016). Lord Howe Island Rodent Eradication Project – Public Environment Report.

Major, R., Ewart, K., Portelli, D., King, A., Tsang, L., O’Dwyer, T., Carlile, N., Haselden, C., Bower, H., & Alquezar-Planas, D. (2021). Islands within islands: genetic structuring at small spatial scales has implications for long-term persistence of a threatened species. Animal Conservation, 24(1), 95–107.

McAllan, I. A., Curtis, B. R., Hutton, I., & Cooper, R. M. (2004). The birds of the Lord Howe Island Group: A review of records. Australian Field Ornithology, 21(Supplement), 1–82.

McCartney-Melstad, E., Gidiş, M., & Shaffer, H. B. (2018). Population genomic data reveal extreme geographic subdivision and novel conservation actions for the declining foothill yellow-legged frog. Heredity, 121(2), 112–125. 10.1038/s41437-018-0097-7

McDougall, I., Embleton, B., & Stone, D. (1981). Origin and evolution of Lord Howe Island, southwest Pacific Ocean. Journal of the Geological Society of Australia, 28(1-2), 155–176.

Meirmans, P. G. (2020). genodive version 3.0: Easy-to-use software for the analysis of genetic data of diploids and polyploids. Molecular Ecology Resources, 20(4), 1126–1131.

Minh, B. Q., Schmidt, H. A., Chernomor, O., Schrempf, D., Woodhams, M. D., von Haeseler, A., & Lanfear, R. (2020). IQ-TREE 2: New models and efficient methods for phylogenetic inference in the genomic era. Molecular Biology and Evolution, 37(5), 1530–1534. 10.1093/molbev/msaa015

Moritz, C. (1994). Defining ‘evolutionarily significant units’ for conservation. Trends in Ecology & Evolution, 9(10), 373–375.

Müller, T., & Juškauskas, A. (2018). Inbreeding affects personality and fitness of a leaf beetle. Animal Behaviour, 138, 29–37.

New South Wales Scientific Committee. (2004). Lord Howe Island Wood-feeding Cockroach – Endangered Species Listing. NSW Scientific Committee, Hurstville, NSW, Australia.

Ney, G., Frederick, K., & Schul, J. (2018). A post-Pleistocene calibrated mutation rate from insect museum specimens. PLOS Currents, 10, ecurrents.tol.aba557de56be881793261f7e1565cf35.

Papadopoulou, A., & Knowles, L.L. (2015). Species-specific responses to island connectivity cycles: refined models for testing phylogeographic concordance across a Mediterranean Pleistocene Aggregate Island Complex. Molecular Ecology, 24, 4252–4268. 10.1111/mec.13305

Papadopoulou, A., & Knowles, L. L. (2017). Linking micro- and macroevolutionary perspectives to evaluate the role of Quaternary sea-level oscillations in island diversification. Evolution, 71(12), 2901–2917. 10.1111/evo.13384

Peel, D., Waples, R. S., Macbeth, G., Do, C., & Ovenden, J. R. (2013). Accounting for missing data in the estimation of contemporary genetic effective population size (Ne). Molecular Ecology Resources, 13(2), 243–253.

Pekkala, N., E. Knott, K., Kotiaho, J. S., Nissinen, K., & Puurtinen, M. (2012). The benefits of interpopulation hybridization diminish with increasing divergence of small populations. Journal of Evolutionary Biology, 25(11), 2181–2193. 10.1111/j.1420-9101.2012.02594.x

Pembleton, L. W., Cogan, N. O. I., & Forster, J. W. (2013). StAMPP: an R package for calculation of genetic differentiation and structure of mixed-ploidy level populations. Molecular Ecology Resources, 13(5), 946–952. 10.1111/1755-0998.12129

Percy, D. M., Page, R. D. M., & Cronk, Q. C. B. (2004). Plant–insect interactions: Double-dating associated insect and plant lineages reveals asynchronous radiations. Systematic Biology, 53(1), 120–127. 10.1080/10635150490264996

Priddel, D., & Wheeler, R. (2014). *New South Wales: state of the islands*. NSW Office of Environment and Heritage, Hurstville, NSW, Australia.

Pritchard, J. K., Stephens, M., & Donnelly, P. (2000). Inference of population structure using multilocus genotype data. Genetics, 155(2), 945–959.

Ralls, K., Ballou, J. D., Dudash, M. R., Eldridge, M. D. B., Fenster, C. B., Lacy, R. C., Sunnucks, P., & Frankham, R. (2018). Call for a paradigm shift in the genetic management of fragmented populations. Conservation Letters, 11(2), e12412. 10.1111/conl.12412

Rambaut, A., Drummond, A. J., Xie, D., Baele, G., & Suchard, M. A. (2018). Posterior summarization in Bayesian phylogenetics using Tracer 1.7. Systematic Biology, 67(5), 901–904. 10.1093/sysbio/syy032

Reid, C. A., Hutton, I., Reid, S. T. C. A., Hutton, I., & Thompson, S. (2020). The citizen scientist survey of large Coleoptera on Lord Howe Island. Technical Reports of the Australian Museum Online, 31, 1–15. 10.3853/j.1835-4211.31.2020.1736

Rijsdijk, K. F., Hengl, T., Norder, S. J., Otto, R., Emerson, B. C., Ávila, S. P., López, H., van Loon, E. E., Tjørve, E., & Fernández-Palacios, J. M. (2014). Quantifying surface-area changes of volcanic islands driven by Pleistocene sea-level cycles: biogeographical implications for the Macaronesian archipelagos. Journal of Biogeography, 41(7), 1242–1254. 10.1111/jbi.12336

Robinson, J. A., Brown, C., Kim, B. Y., Lohmueller, K. E., & Wayne, R. K. (2018). Purging of strongly deleterious mutations explains long-term persistence and absence of inbreeding depression in island foxes. Current Biology, 28(21), 3487–3494. e3484.

Rogers, A., Flanigan, M., Nebel, O., Nebel-Jacobsen, Y., Wang, X., Arculus, R. J., Miller, L., Smith, I., Mather, B. R., & Kendrick, M. (2023). The isotopic origin of Lord Howe Island reveals secondary mantle plume twinning in the Tasman Sea. Chemical Geology, 622, 121374.

Rose, H. (2003). Research Report on Panesthia lata, Blaberid cockroach 25-29 March 2003.

Sheringham, P., Richards, P., Gilmour, P., & Kemmerer, E. (2016). A Systematic Flora Survey, Floristic Classification and High-Resolution Vegetation Map of Lord Howe Island. Lord Howe Island Board. 10.13140/RG.2.2.17814.45123

Spratt, R. M., & Lisiecki, L. E. (2016). A Late Pleistocene sea level stack. Climate of the Past, 12(4), 1079–1092. 10.5194/cp-12-1079-2016

Sturm, A. B., Eckert, R. J., Méndez, J. G., González-Díaz, P., & Voss, J. D. (2020). Population genetic structure of the great star coral, *Montastraea cavernosa*, across the Cuban archipelago with comparisons between microsatellite and SNP markers. Scientific Reports, 10(1), 15432.

Suchard, M. A., Lemey, P., Baele, G., Ayres, D. L., Drummond, A. J., & Rambaut, A. (2018). Bayesian phylogenetic and phylodynamic data integration using BEAST 1.10. Virus Evolution, 4(1), vey016.

Tallmon, D. A., Luikart, G., & Waples, R. S. (2004). The alluring simplicity and complex reality of genetic rescue. Trends in Ecology & Evolution, 19(9), 489–496.

Team, R. (2020). RStudio: Integrated Development for R. In RStudio, PBC. http://www.rstudio.com/

Walker, F. (1868). Catalogue of the specimens of Blattariæ in the collection of the British Museum. Printed for the Trustees of the British Museum. https://www.biodiversitylibrary.org/item/35169

Waples, R. S. (2006). A bias correction for estimates of effective population size based on linkage disequilibrium at unlinked gene loci. Conservation Genetics, 7(2), 167–184.

Weeks, A. R., Heinze, D., Perrin, L., Stoklosa, J., Hoffmann, A. A., van Rooyen, A., Kelly, T., & Mansergh, I. (2017). Genetic rescue increases fitness and aids rapid recovery of an endangered marsupial population. Nature Communications, 8(1), 1071. 10.1038/s41467-017-01182-3

Weir, B. S., & Cockerham, C. C. (1984). Estimating *F*-statistics for the analysis of population structure. Evolution, 38, 1358–1370.

Whiteley, A. R., Fitzpatrick, S. W., Funk, W. C., & Tallmon, D. A. (2015). Genetic rescue to the rescue. Trends in Ecology & Evolution, 30(1), 42–49. 10.1016/j.tree.2014.10.009

Wilkinson, I., & Priddel, D. (2011). Rodent eradication on Lord Howe Island: challenges posed by people, livestock, and threatened endemics. In: Island Invasives: Eradication and Management (eds Veitch, C. R., Clout, M. N., Towns, D. R.) IUCN: Gland, Switzerland, 508–514.

Younger, J. L., Clucas, G. V., Kao, D., Rogers, A. D., Gharbi, K., Hart, T., & Miller, K. J. (2017). The challenges of detecting subtle population structure and its importance for the conservation of emperor penguins. Molecular Ecology, 26(15), 3883–3897.

Zheng, X., & Zheng, M. X. (2013). Package ‘SNPRelate’. A package for parallel computing toolset for relatedness and principal component analysis of SNP data.

Zink, R. M. (2004). The role of subspecies in obscuring avian biological diversity and misleading conservation policy. Proceedings of the Royal Society B, 271(1539), 561–564.

Zwick, P., & Zwick, A. (2023). Revision of the African *Neoperla* Needham, 1905 (Plecoptera: Perlidae: Perlinae) based on morphological and molecular data. Zootaxa, 5316(1), 1–194.

