## Supplementary Information for "Pleistocene sea-level fluctuation shapes archipelago-wide population structure in the Endangered Lord Howe Island cockroach *Panesthia lata*"

**Supplementary material**

**Supplementary Table S1.** Samples of *Panesthia lata* and outgroup taxa used in the study. Abbreviations: Australian Museum (AM), Macleay Museum (MM), H.A. Rose private collection (HARPC), single-nucleotide polymorphism (SNP). Missing data quantified prior to filtering.

| **Sample ID / species** | **Locality** | **Institution** | **Year collected** | **SNPs genotyped** | **SNP missing data (%)** | **Mitogenome accession number** |
| --- | --- | --- | --- | --- | --- | --- |
| K.487941 | Ball’s Pyramid | AM | 1969 | Y | 88.5 | PQ231654 |
| K.487942 | Ball’s Pyramid | AM | 1969 | Y | 84.7 | - |
| PL17 | Blackburn Island | HARPC | 2022 | Y | 21.9 | PQ231560 |
| PL3 | Blackburn Island | HARPC | 2022 | Y | 21.2 | PQ231639 |
| PL33 | Blackburn Island | HARPC | 2022 | Y | 19.0 | - |
| PL34 | Blackburn Island | HARPC | 2022 | Y | 18.2 | - |
| PL35 | Blackburn Island | HARPC | 2022 | Y | 18.4 | - |
| PL36 | Blackburn Island | HARPC | 2022 | Y | 18.4 | - |
| PL38 | Blackburn Island | HARPC | 2022 | Y | 19.4 | PQ231639 |
| PL4 | Blackburn Island | HARPC | 2022 | Y | 21.0 | - |
| PL41 | Blackburn Island | HARPC | 2022 | Y | 19.2 | PQ231562 |
| PL42 | Blackburn Island | HARPC | 2022 | Y | 18.3 | - |
| PL45 | Blackburn Island | HARPC | 2022 | Y | 18.7 | - |
| PL48 | Blackburn Island | HARPC | 2022 | Y | 21.0 | - |
| PL57 | Blackburn Island | HARPC | 2022 | Y | 18.8 | - |
| PL6 | Blackburn Island | HARPC | 2022 | Y | 19.2 | PQ231637 |
| PL7 | Blackburn Island | HARPC | 2022 | Y | 21.6 | - |
| K.487917 | Lord Howe Island | AM | 1869–1950 | N | - | - |
| K.487918 | Lord Howe Island | AM | 1869–1950 | N | - | - |
| K.487919 | Lord Howe Island | AM | 1869–1950 | N | - | - |
| K.487920 | Lord Howe Island | AM | 1869–1950 | N | - | - |
| K.487921 | Lord Howe Island | AM | 1869–1950 | N | - | PQ231559 |
| K.487922 | Lord Howe Island | AM | 1869–1950 | N | - | - |
| K.487923 | Lord Howe Island | AM | 1869–1950 | N | - | - |
| K.487924 | Lord Howe Island | AM | 1869–1950 | N | - | - |
| K.487925 | Lord Howe Island | AM | 1869–1950 | N | - | - |
| K.487926 | Lord Howe Island | AM | 1869–1950 | N | - | - |
| K.487927 | Lord Howe Island | AM | 1869–1950 | N | - | - |
| K.487928 | Lord Howe Island | AM | 1869–1950 | N | - | - |
| K.487929 | Lord Howe Island | AM | 1869–1950 | N | - | - |
| K.487930 | Lord Howe Island | AM | 1869–1950 | N | - | PQ231638 |
| K.487931 | Lord Howe Island | AM | 1869–1950 | N | - | - |
| K.487932 | Lord Howe Island | AM | 1869–1950 | N | - | - |
| K.487934 | Lord Howe Island | AM | 1869–1950 | N | - | - |
| K.487935 | Lord Howe Island | AM | 1869–1950 | N | - | - |
| K.487936 | Lord Howe Island | AM | 1869–1950 | N | - | - |
| K.487937 | Lord Howe Island | AM | 1869–1950 | N | - | - |
| NHEN.66552 | Lord Howe Island | MM | 1869–1950 | N | - | PQ248021 |
| NHEN.66559 | Lord Howe Island | MM | 1869–1950 | N | - | PQ248022 |
| NHEN.66554 | Lord Howe Island | MM | 1869–1950 | N | - | PQ248023 |
| NHEN.66555 | Lord Howe Island | MM | 1869–1950 | N | - | PQ248024 |
| PL1 | North Bay | HARPC | 2022 | Y | 22.9 | PQ231567 |
| PL18 | North Bay | HARPC | 2022 | Y | 21.7 | - |
| PL19 | North Bay | HARPC | 2022 | Y | 27.5 | - |
| PL2 | North Bay | HARPC | 2022 | Y | 25.7 | - |
| PL26 | North Bay | HARPC | 2022 | Y | 23.7 | - |
| PL27 | North Bay | HARPC | 2022 | Y | 21.5 | PQ231563 |
| PL28 | North Bay | HARPC | 2022 | Y | 22.3 | PQ231564 |
| PL29 | North Bay | HARPC | 2022 | Y | 21.8 | - |
| PL30 | North Bay | HARPC | 2022 | Y | 21.5 | - |
| PL5 | North Bay | HARPC | 2022 | Y | 21.2 | - |
| PL21 | Roach Island | HARPC | 2022 | Y | 25.3 | - |
| PL22 | Roach Island | HARPC | 2022 | Y | 22.0 | - |
| PL23 | Roach Island | HARPC | 2022 | Y | 21.8 | - |
| PL24 | Roach Island | HARPC | 2022 | Y | 22.5 | - |
| PL25 | Roach Island | HARPC | 2022 | Y | 22.9 | - |
| PL66 | Roach Island | HARPC | 2022 | Y | 35.7 | - |
| K.383796 | Roach Island | AM | 2000 | Y | 27.3 | PQ231565 |
| K.383797 | Roach Island | AM | 2000 | Y | 88.5 | - |
| K.383798 | Roach Island | AM | 2000 | Y | 49.0 | PQ231569 |
| K.383799 | Roach Island | AM | 2000 | Y | 41.2 | PQ231636 |
| K.383800 | Roach Island | AM | 2000 | Y | 82.9 | PQ231566 |
| K.383801 | Roach Island | AM | 2000 | Y | 23.8 | - |
| PL10 | Roach Island | HARPC | 2003 | Y | 29.5 | - |
| PL11 | Roach Island | HARPC | 2003 | Y | 25.8 | - |
| PL12 | Roach Island | HARPC | 2003 | Y | 22.8 | - |
| PL13 | Roach Island | HARPC | 2003 | Y | 22.7 | - |
| PL14 | Roach Island | HARPC | 2003 | Y | 26.3 | - |
| PL15 | Roach Island | HARPC | 2003 | Y | 26.3 | - |
| PL16 | Roach Island | HARPC | 2003 | Y | 24.2 | - |
| PL20 | Roach Island | HARPC | 2003 | Y | 27.5 | - |
| PL8 | Roach Island | HARPC | 2003 | Y | 34.0 | - |
| PL9 | Roach Island | HARPC | 2003 | Y | 33.4 | - |
| *Panesthia cribrata* | - | - | - | - | - | PQ231600 |
| *Panesthia matthewsi* | - | - | - | - | - | PQ231605 |

**Supplementary Table S2.** Best-fitting substitution models for each partition in the mitogenomic data sets, as selected by ModelFinder (Kalyaanamoorthy et al., 2017). Abbreviations: protein-coding genes (PCG), ribosomal RNA (rRNA), transfer RNA (tRNA).

| **Partition** | **Length (bp)** | **Substitution model** |
| --- | --- | --- |
| *CO1* | 1533 | HKY+F+R2 |
| PCG codon 1 | 3187 | TIM2+F+I |
| PCG codon 2 | 3187 | HKY+F+I |
| PCG codon 3 | 3187 | TIM+F+G4 |
| rRNA | 2130 | TIM3+F+I |
| tRNA | 1367 | HKY+F+I |

**Supplementary Table S3.** Pairwise *F*_ST_ values (95% confidence interval) between 3 populations of *Panesthia lata* based on 9,283 nuclear SNPs from 36 unrelated individuals (* denotes values significantly different from 0). Closely related individuals were identified and removed based on kinship ≥ 0.125, as calculated in *SNPRelate* (Zheng & Zheng, 2013).

|  | **Roach Island** | **Blackburn Island** |
| --- | --- | --- |
| **Blackburn Island** | 0.234* (0.225–0.244) | - |
| **North Bay** | 0.229* (0.216–0.242) | 0.203* (0.189–0.217) |

**Supplementary Table S4.** Summary of genetic diversity statistics across three populations of *Panesthia lata*, including only Roach Island individuals from a single sampling batch collected in December 2003. *N* = sample size, *A*_R_ = mean rarefied allelic richness, *H*_O_ = mean observed heterozygosity, SE = standard error, *H*_O_ = mean expected heterozygosity, *F*_IS_ = inbreeding coefficient (* denotes value significantly different from 0), 95% CI = 95% lower and upper confidence interval, *N*_e_ = effective population size. All values except *N*_e_ based on 9,729 SNPs, N_e_ values based on 12,027 SNPs.

| **Population** | ***N*** | ***A*_R_** | ***H*_O_ (SE)** | ***H*_E_ (SE)** | ***F*_IS_ (95% CI)** | ***N*_e_ (95% CI)** |
| --- | --- | --- | --- | --- | --- | --- |
| **Roach Island** | 11 | 1.491 | 0.116 (0.049) | 0.192 (0.060) | 0.388* (0.378–0.400) | 258.2 (5.5–Inf.) |
| **Blackburn Island** | 15 | 1.439 | 0.125 (0.039) | 0.167 (0.044) | 0.258* (0.249–0.267) | 1247.4 (340.8–Inf.) |
| **North Bay** | 10 | 1.410 | 0.121 (0.054) | 0.165 (0.061) | 0.275* (0.264–0.286) | 47.1 (28.4–118.6) |

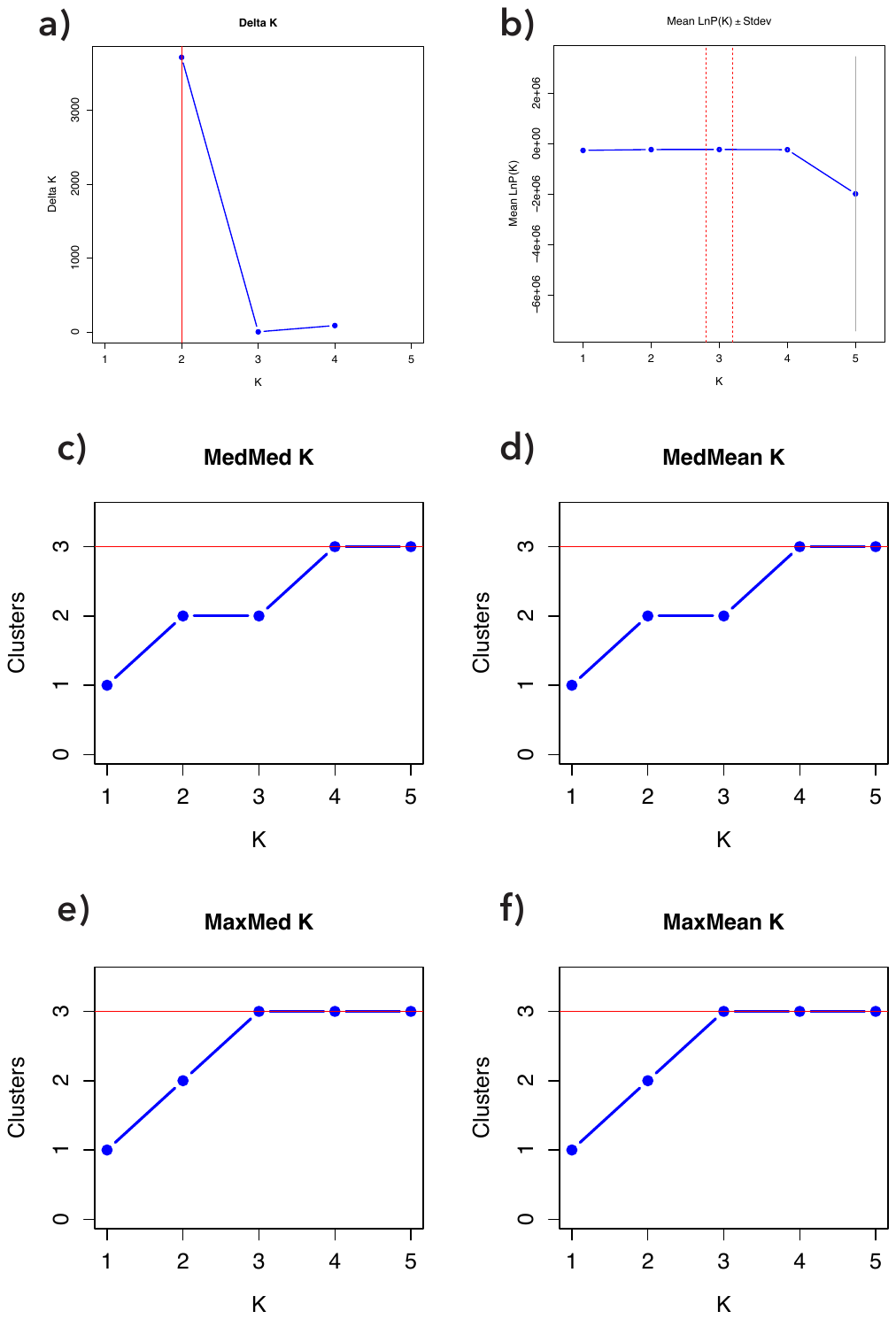
**Supplementary Figure S1.** Estimated values of *K* based on 8,012 nuclear SNPs from 44 individuals of *Panesthia lata*. The metrics used were: **a)** the Delta K metric (Evanno et al., 2005); **b)** the log-likelihood of posterior probability metric (Pritchard et al., 2000); and the four metrics presented by (Puechmaille, 2016): **c)** the median of medians, **d)** the median of means, **e)** the maximum of medians, and **f)** the maximum of means.

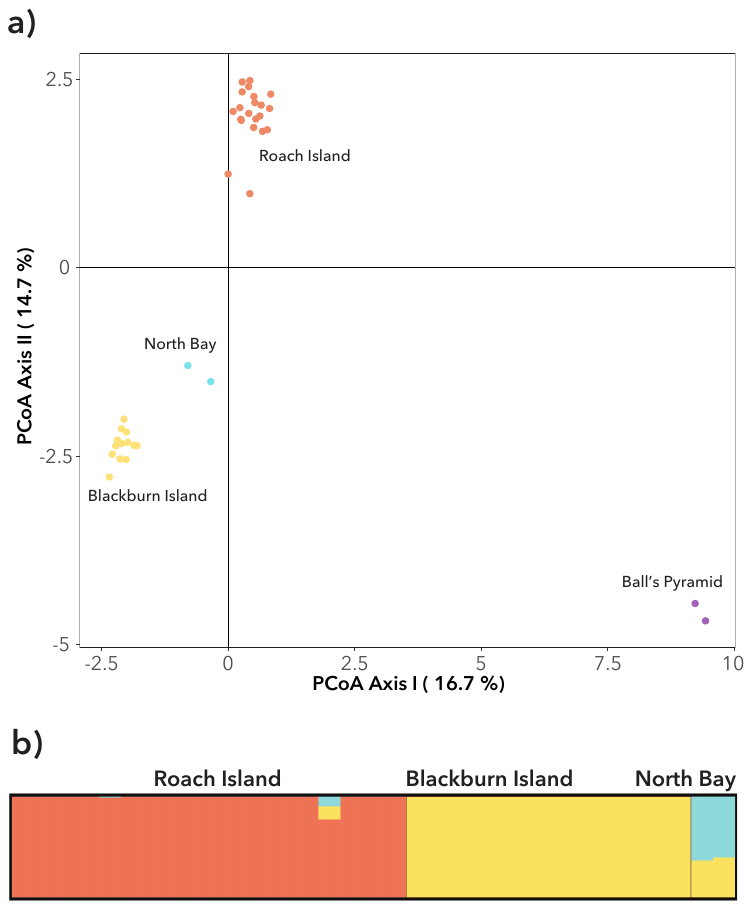
**Supplementary Figure S2.** **a)** Principal coordinates analysis (PCoA) plot of 38 unrelated individuals of *Panesthia lata* using 544 nuclear single-nucleotide polymorphisms (SNPs). **b)** STRUCTURE plots for 33 unrelated individuals using 9,232 SNPs, when *K* = 3. Individuals from Ball’s Pyramid were omitted from this data set. Closely related individuals were identified and removed based on kinship ≥ 0.125, as calculated in *SNPRelate* (Zheng & Zheng, 2013)

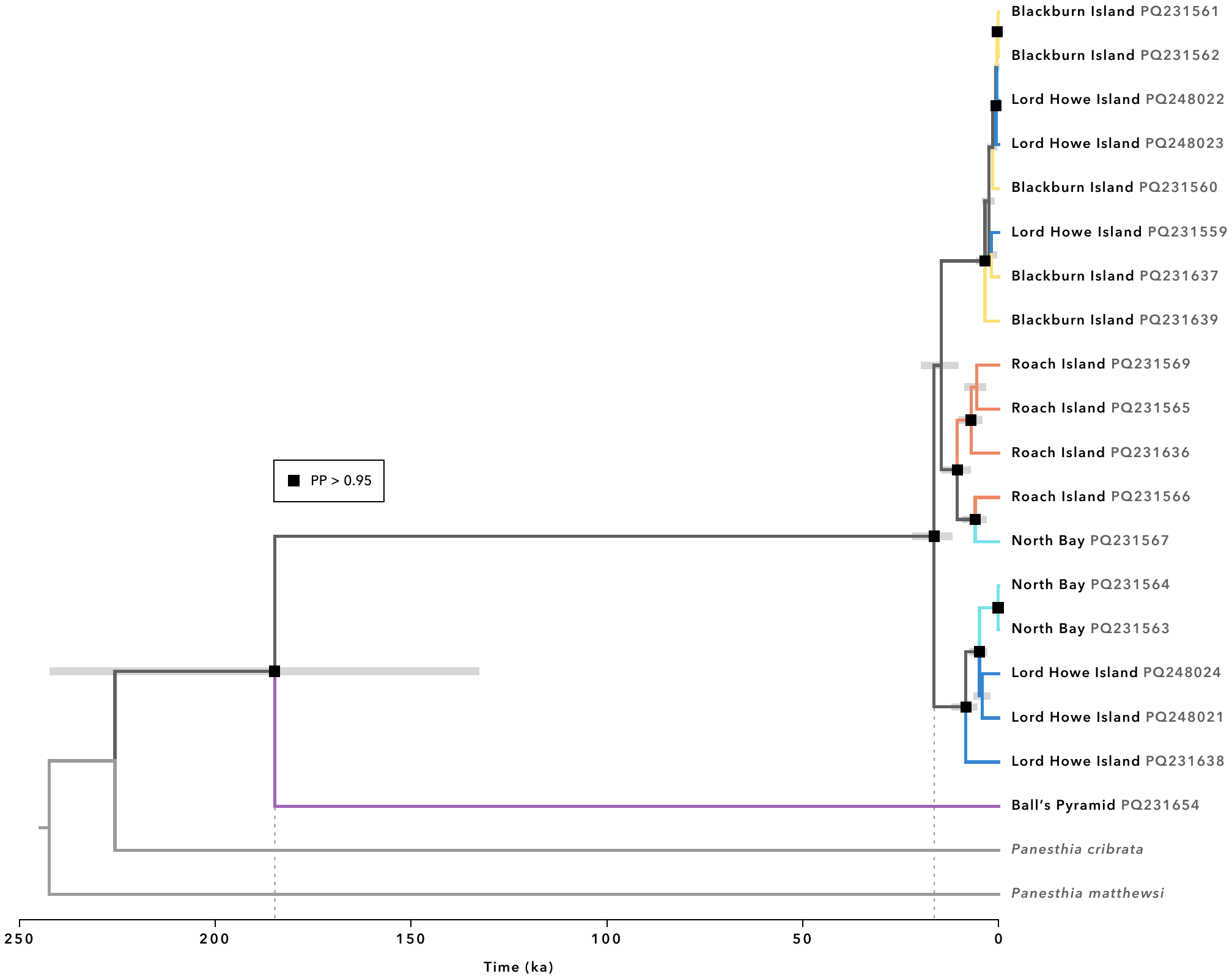
**Supplementary Figure S3.** Dated phylogeny of *Panesthia lata* inferred from complete mitochondrial genomes in BEAST. The figure depicts our minimum age estimate, which was derived by specifying an informative prior on the substitution rate of *CO1* (Ney et al., 2018). PP: posterior probability. Numbers on labels indicate GenBank accession numbers.

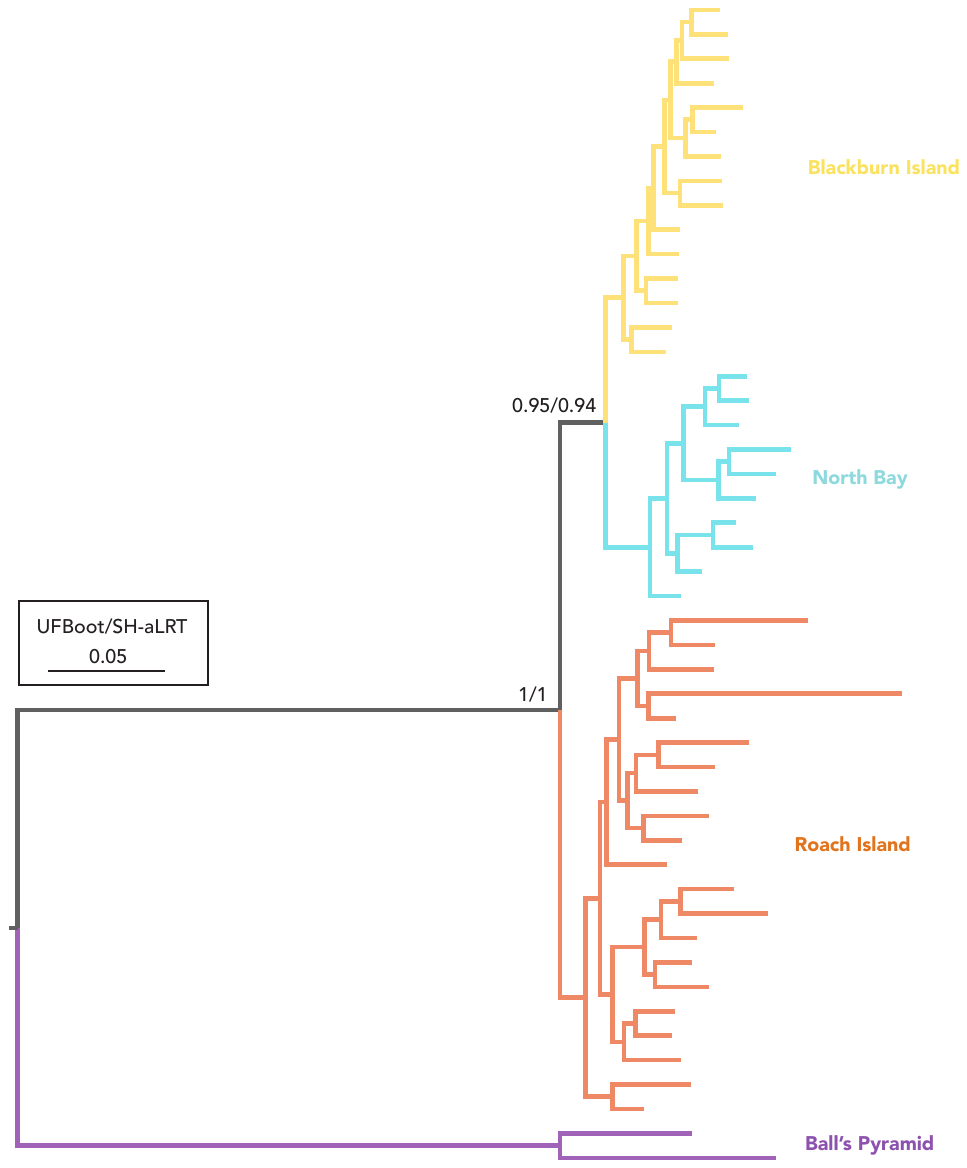

**Supplementary Figure S4.** Maximum-likelihood phylogeny of *Panesthia lata* inferred from 1,315 nuclear SNPs in IQTREE. UFBoot: ultrafast bootstrap, SH-aLRT: SH-like likelihood-ratio test. The root was placed between samples from Ball’s Pyramid and remaining populations. Node support within populations not shown.

**References**

Evanno, G., Regnaut, S., & Goudet, J. (2005). Detecting the number of clusters of individuals using the software STRUCTURE: a simulation study. *Molecular Ecology*, *14*(8), 2611-2620.

Kalyaanamoorthy, S., Minh, B. Q., Wong, T. K., Von Haeseler, A., & Jermiin, L. S. (2017). ModelFinder: fast model selection for accurate phylogenetic estimates. *Nature Methods*, *14*(6), 587-589.

Ney, G., Frederick, K., & Schul, J. (2018). A post-pleistocene calibrated mutation rate from insect museum specimens. *PLoS Currents*, *10*.

Pritchard, J. K., Stephens, M., & Donnelly, P. (2000). Inference of population structure using multilocus genotype data. *Genetics*, *155*(2), 945-959.

Puechmaille, S. J. (2016). The program structure does not reliably recover the correct population structure when sampling is uneven: subsampling and new estimators alleviate the problem. *Molecular ecology resources*, *16*(3), 608-627.

Zheng, X., & Zheng, M. X. (2013). Package ‘SNPRelate’. *A package for parallel computing toolset for relatedness and principal component analysis of SNP data*.
